# Disruption of IRE1α through its Kinase Domain Attenuates Multiple Myeloma

**DOI:** 10.1101/495242

**Authors:** Jonathan M Harnoss, Adrien Le Thomas, Scot A Marsters, David A Lawrence, Min Lu, Yung-Chia Ariel Chen, Jing Qing, Klara Totpal, David Kan, Ehud Segal, Heidi Ackerly Wallweber, Weiru Wang, Kevin Clark, Susan Kaufman, Maureen Beresini, Wendy Sandoval, Maria Lorenzo, Jiansheng Wu, Justin Ly, Tom De Bruyn, Amy Heidersbach, Benjamin Haley, Alvin Gogineni, Robby Weimer, Dong Lee, Marie-Gabrielle Braun, Joachim Rudolph, Michael J VanWyngarden, Daniel W Sherbenou, Patricia Gomez-Bougie, Martine Amiot, Diego Acosta-Alvear, Peter Walter, Avi Ashkenazi

**Affiliations:** Cancer Immunology, Genentech, Inc., 1 DNA Way, South San Francisco, CA 94080 USA; Translational Oncology, 1 DNA Way, South San Francisco, CA 94080 USA; Structural Biology, 1 DNA Way, South San Francisco, CA 94080 USA; Biochemical and Cellular Pharmacology, Genentech, Inc. 1 DNA Way, South San Francisco, CA 94080 USA; Microchemistry, Proteomics and Lipidomics, Genentech, Inc. 1 DNA Way, South San Francisco, CA 94080 USA; Protein Chemistry, Genentech, Inc. 1 DNA Way, South San Francisco, CA 94080 USA; Drug Metabolism and Pharmacokinetics, Genentech, Inc. 1 DNA Way, South San Francisco, CA 94080 USA; Molecular Biology, Genentech, Inc. 1 DNA Way, South San Francisco, CA 94080 USA; Biomolecular Imaging, Genentech, Inc. 1 DNA Way, South San Francisco, CA 94080 USA; Safety Assessment, Genentech, Inc. 1 DNA Way, South San Francisco, CA 94080 USA; Discovery Chemistry, Genentech, Inc. 1 DNA Way, South San Francisco, CA 94080 USA; Division of Hematology, Department of Medicine, University of Colorado Cancer Center, University of Colorado Anschutz Medical Campus, Aurora CO 80045 USA; CRCINA, INSERM, CNRS, Université d’Angers, Université de Nantes, Nantes, BP 70721, France; Service d’Hématologie Clinique, Unité d’Investigation Clinique, CHU, Nantes, BP 70721 France; Department of Biochemistry and Biophysics, University of California San Francisco, San Francisco, CA 94143 USA; Howard Hughes Medical Institute, University of California San Francisco, San Francisco, CA 94143 USA

**Author notes:** Agios Pharmaceuticals, 88 Sidney Street, Cambridge, MA 02139 USA. Revolution Medicines, 700 Saginaw Dr., Redwood City, CA 94063. Department of Molecular, Cellular and Developmental Biology, University of California Santa Barbara, Santa Barbara, CA 93106 USA.

**Keywords:** Multiple myeloma, endoplasmic reticulum stress, unfolded protein response, inositol requiring enzyme 1, kinase inhibitors

## Abstract

Multiple myeloma (MM) arises from malignant immunoglobulin-secreting plasma cells and remains an incurable, often lethal disease despite recent therapeutic advances. The unfolded-protein response sensor IRE1α supports protein secretion by deploying a kinase-endoribonuclease module to activate the transcription factor XBP1s. MM cells may coopt the IRE1α-XBP1s pathway; however, the validity of IRE1α as a potential MM therapeutic target is controversial. Here we show that genetic disruption of IRE1α or XBP1s, or pharmacologic IRE1α kinase inhibition, attenuated subcutaneous or orthometastatic growth of MM tumors in mice, and augmented efficacy of two well-established frontline antimyeloma agents, bortezomib or lenalidomide. Mechanistically, IRE1α perturbation inhibited expression of key components of the ER-associated degradation machinery, as well as cytokines and chemokines known to promote MM growth. Selective IRE1α kinase inhibition reduced viability of CD138^+^ plasma cells while sparing CD138^−^ cells from bone marrow of newly diagnosed MM patients or patients whose disease relapsed after 1 - 4 lines of treatment in both US- and EU-based cohorts. IRE1α inhibition preserved survival and glucose-induced insulin secretion by pancreatic microislets. Together, these results establish a strong therapeutic rationale for targeting IRE1α with kinase-based small-molecule inhibitors in MM.

**Significance statement:** Multiple myeloma (MM) is a lethal malignancy of plasma cells. MM cells have an expanded endoplasmic reticulum (ER) that is constantly under stress due to immunoglobulin hyperproduction. The ER-resident sensor IRE1α mitigates ER stress by expanding the ER’s protein-folding capacity while supporting proteasomal degradation of misfolded ER proteins. IRE1α elaborates these functions by deploying its cytoplasmic kinase-RNase module to activate the transcription factor XBP1s. The validity of IRE1α as a potential therapeutic target in MM has been questioned. Using genetic and pharmacologic disruption in vitro and in vivo, we demonstrate that the IRE1α-XBP1s pathway plays a critical role in MM growth. We further show that IRE1α’s kinase domain is an effective and safe potential small-molecule target for MM therapy.

## Introduction

MM is the second most common human hematologic cancer. It carries a lifetime risk of 0.7% and occurs mainly in older individuals. MM is caused by bone marrow infiltration of malignant, monoclonal immunoglobulin (Ig-)-secreting plasma cells (1). Despite significant therapeutic advances—including proteasome inhibitors (PIs), immunomodulatory agents (IMiDs), and anti-CD38 antibodies—MM remains mainly incurable, with acquired resistance to all available agents, and 5-year survival of 49% (2). Hence, considering the rapid growth of the aging population in many countries, there is an urgent need for development of novel MM therapies.

The endoplasmic reticulum (ER) assures the precise folding of newly synthesized transmembrane and secreted proteins. Upon elevations in cellular demand for protein secretion— for example, when mature B cells differentiate into Ig-secreting plasma cells—insufficient ER capacity causes accumulation of unfolded proteins (UPs) in the ER lumen. This activates a rheostatic sensing-signaling network—dubbed the unfolded protein response (UPR)—to orchestrate ER adaptation and reestablish cellular homeostasis (3-6). The mammalian UPR employs three pivotal ER-resident transmembrane sensors: inositol-requiring enzyme 1 alpha (IRE1α), protein kinase-like endoplasmic reticulum kinase (PERK), and activating transcription factor-6 (ATF6). Detection of excess UPs by the ER lumenal domain of each sensor engages its cytoplasmic moiety to expand the ER and its protein folding and degradative capacities, thus alleviating ER stress. If this corrective response fails and stress becomes overwhelming, the UPR can trigger apoptosis (7). Conserved from yeast to primates, IRE1α harbors lumenal, transmembrane and cytosolic regions: The cytoplasmic part contains two tandem catalytic modules: A serine/threonine kinase and an endoribonuclease (RNase) (8, 9). IRE1α activation involves homodimerization, trans-autophosphorylation of the cytoplasmic kinase domain, and RNase activation (9-12). The RNase triggers non-conventional splicing of the mRNA encoding X-box protein 1 (XBP1) by removing a 26-nucleotide intron to produce spliced XBP1 (XBP1s) (13). XBP1s encodes a potent transcription factor that activates multiple genes encoding chaperones, disulfide isomerases and lipid synthases, thereby facilitating biochemical and biophysical expansion of the ER (14-16). Additional XBP1s gene-targets encode components of the ER-associated degradation (ERAD) machinery, which promotes retro-translocation of UPs into the cytoplasm, followed by their ubiquitination and proteasomal disposal (14, 17). An additional activity of IRE1α’s RNase—termed regulated IRE1-dependent RNA decay (RIDD)—cleaves ER-associated mRNAs, temporarily abating translational load (18, 19) and suppressing apoptosis (20, 21), which allows time for cellular adaptation.

Because plasma-cell differentiation requires IRE1α and XBP1s (22-24), and because cancer cells often coopt normal stress-response pathways to support malignant growth in hostile microenvironments (25), it has been proposed that the IRE1α-XBP1s pathway may represent a therapeutically useful vulnerability in MM cells (26-28). Supporting this hypothesis, transgenic expression of XBP1s in B cells drove a MM-like disease in mice (29) and high XBP1s levels correlated with worse prognosis in MM patients (30). XBP1s depletion by shRNA attenuated growth of certain MM cell lines *in vitro*, and small-molecule inhibition of IRE1α’s RNase activity with salicylaldehyde compounds attenuated human MM xenograft growth in mice (31, 32). Standard-of-care agents such as PIs are effective in MM therapy likely because their inhibition of the 26S proteasome creates a backlog of ERAD substrates that cannot be efficiently degraded, thereby exacerbating ER stress (33). However, conflicting evidence suggests either positive (33, 34) or negative (35) correlations between XBP1s levels and MM responsiveness to PI therapy. Moreover, although IRE1α kinase inhibition blocked XBP1s production, it failed to attenuate MM cell growth *in vitro* (36). An additional concern is that the selectivity of the highly protein-reactive salicylaldehyde-based IRE1α RNase inhibitors is difficult to ascertain: indeed, recent evidence demonstrates that one of these compounds causes oxidative stress due to off-target activity (37). Thus, whether IRE1α can be targeted effectively and safely to inhibit malignant MM growth remains controversial. In addition, because XBP1s depletion drives hyper-phosphorylation of IRE1α (20, 38), alternative, XBP1s-independent IRE1α functions— for example, activation of c-Jun N-terminal kinase (JNK) (39)—also may impact MM cells. Another key question that remains unanswered in this context is whether IRE1α can be targeted effectively and safely via its kinase domain to inhibit MM.

Our present results demonstrate that the IRE1α-XBP1s pathway plays a critical role in supporting MM cell growth *in vitro* in 3D culture, as well as *in vivo* in subcutaneous and orthometastatic tumor xenograft settings. Selective small-molecule IRE1α kinase inhibition reduced viability of patient-derived malignant MM cells yet spared accompanying normal cells from bone marrow samples, and also preserved insulin secretion by pancreatic microislets. Together, these findings establish a compelling rationale for developing small-molecule inhibitors targeting IRE1α kinase for MM therapy.

## Results

### Depletion of IRE1α by shRNAs attenuates 3D growth of MM cell lines

Interrogation of the cancer cell line encyclopedia (CCLE) RNAseq dataset (Broad Institute, Cambridge, MA, USA) demonstrated that MM cell lines express higher mRNA levels of IRE1α than all other cancer types (**SI Appendix, Fig. S1*A***). Immunoblot (IB) analysis of 12 human MM cell lines revealed abundant IRE1α protein, often in conjunction with detectable XBP1s protein (**Fig. *1A***), suggesting frequent IRE1α-XBP1s pathway activation in MM cells. To investigate the importance of IRE1α for MM cell growth, we first used a doxycycline-(Dox-) inducible shRNA-based knockdown approach. As expected, Dox-driven anti-IRE1α shRNA expression markedly decreased IRE1α and XBP1s protein levels in KMS11, OPM2 and RPMI8226 MM cells (**Fig. *1B***). Importantly, Dox-induced IRE1α depletion markedly inhibited 3D growth of these three cell lines in the form of single spheroids on ultra-low attachment plates, as evident by fluorescence imaging (**Fig. *1C-H***). IRE1α knockdown also inhibited 3D growth of KMS11 cells in the form of multiple spheroids on matrigel, as determined via an Incucyte™ instrument (**SI Appendix, Fig. S1*B-E***). Of note, similar Dox treatment did not alter growth or viability of the parental KMS11 cells (**SI Appendix**, **Fig. S1*B, D*** and ***F***), or parental OPM2 and RPMI8226 cells (data not shown). In concert with the growth inhibition, IRE1α depletion in KMS11 cells cultured on matrigel led to a substantial and specific loss of viability, as measured by a CellTiterGlo^^®^^ assay (**SI Appendix, Fig. S1*F-H***). Thus, three genetically diverse MM cell lines (40) displayed significant dependence on IRE1α for 3D growth—a modality that more faithfully reflects *in vivo* tumor settings than the conventional 2D culture used in earlier work (36).

**Fig. 1.**
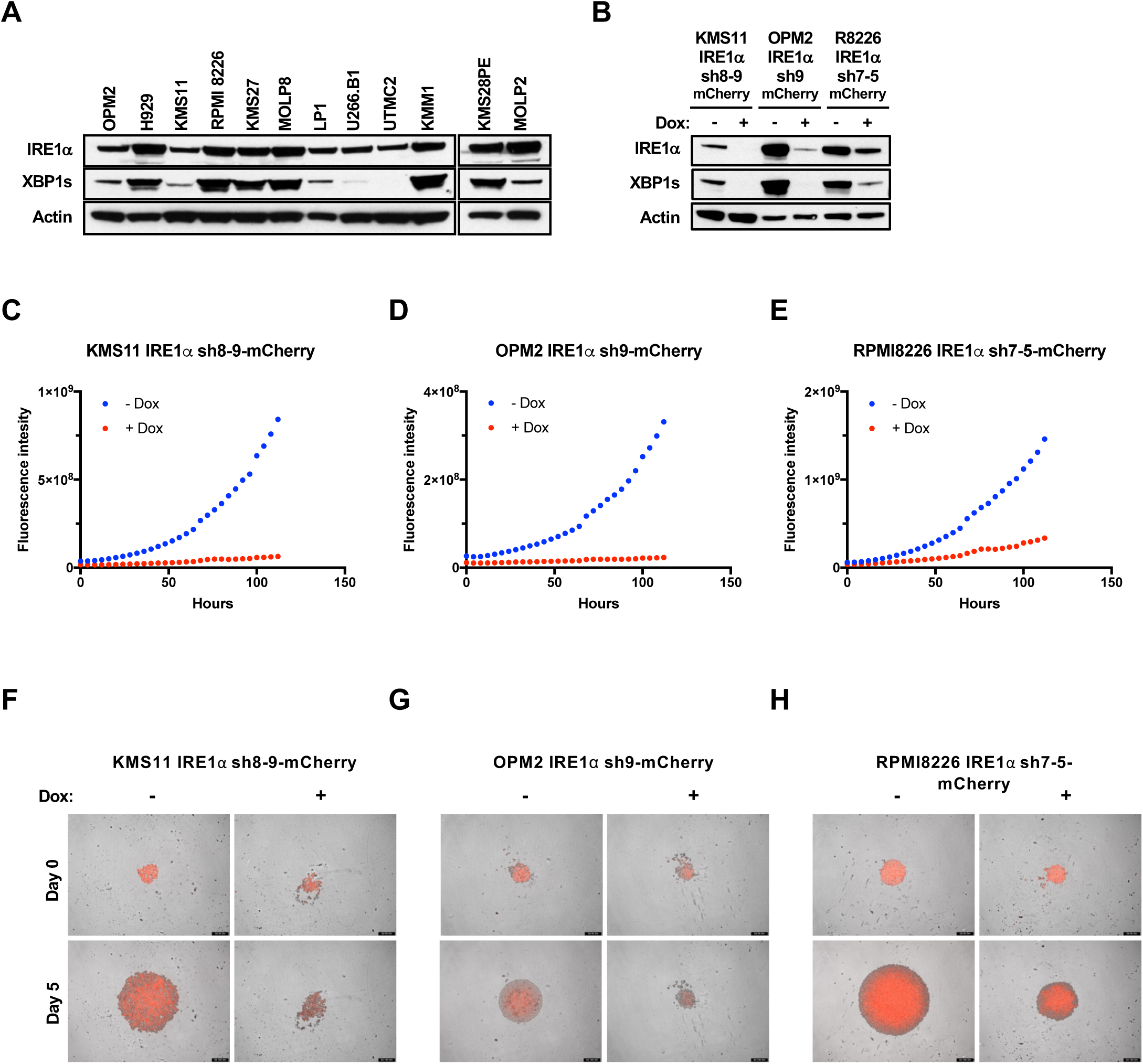
Expression of IRE1α in MM cell lines and effect of its depletion on spheroid 3D growth. **(*A*)** Twelve human MM cell lines were analyzed by immunoblot (IB) for protein levels of IRE1α and XBP1s. (***B***) KMS11, OPM2, and RPMI8226 MM cells were stably transfected with a plasmid encoding Dox-inducible shRNA against IRE1α together with a plasmid encoding mCherry. Cells were incubated in the absence or presence of Dox (0.5 μg/ml) for 3 days, seeded on ultra-low adhesion plates, centrifuged to form single spheroids, and analyzed by IB for indicated proteins (***B***), or for growth based on mCherry fluorescence (***C-E***), or imaged by fluorescence microscopy (***F-H***).

### Genetic disruption of IRE1α or XBP1s attenuates growth of subcutaneous human MM xenografts

We next disrupted the IRE1α gene in KMS11 cells using CRISPR/Cas9 gene editing. In contrast to parental KMS11 cells, three independent IRE1α KO clones showed a complete absence of IRE1α protein and failed to upregulate XBP1s in response to the ER stressor thapsigargin (Tg) (**SI Appendix, Fig. S2*A***). Upon subcutaneous injection into C.B-17 SCID mice, wildtype (WT) KMS11 cells formed readily palpable tumors that reached a mean volume of ~500 mm^3^ by 29 days; in contrast, all three IRE1α KO KMS11 clones failed to sustain appreciable tumor growth (**Fig. 2*A*; SI Appendix, Fig. S2*B***). Thus, subcutaneous establishment and growth of KMS11 MM xenografts in mice requires IRE1α.

**Fig. 2.**
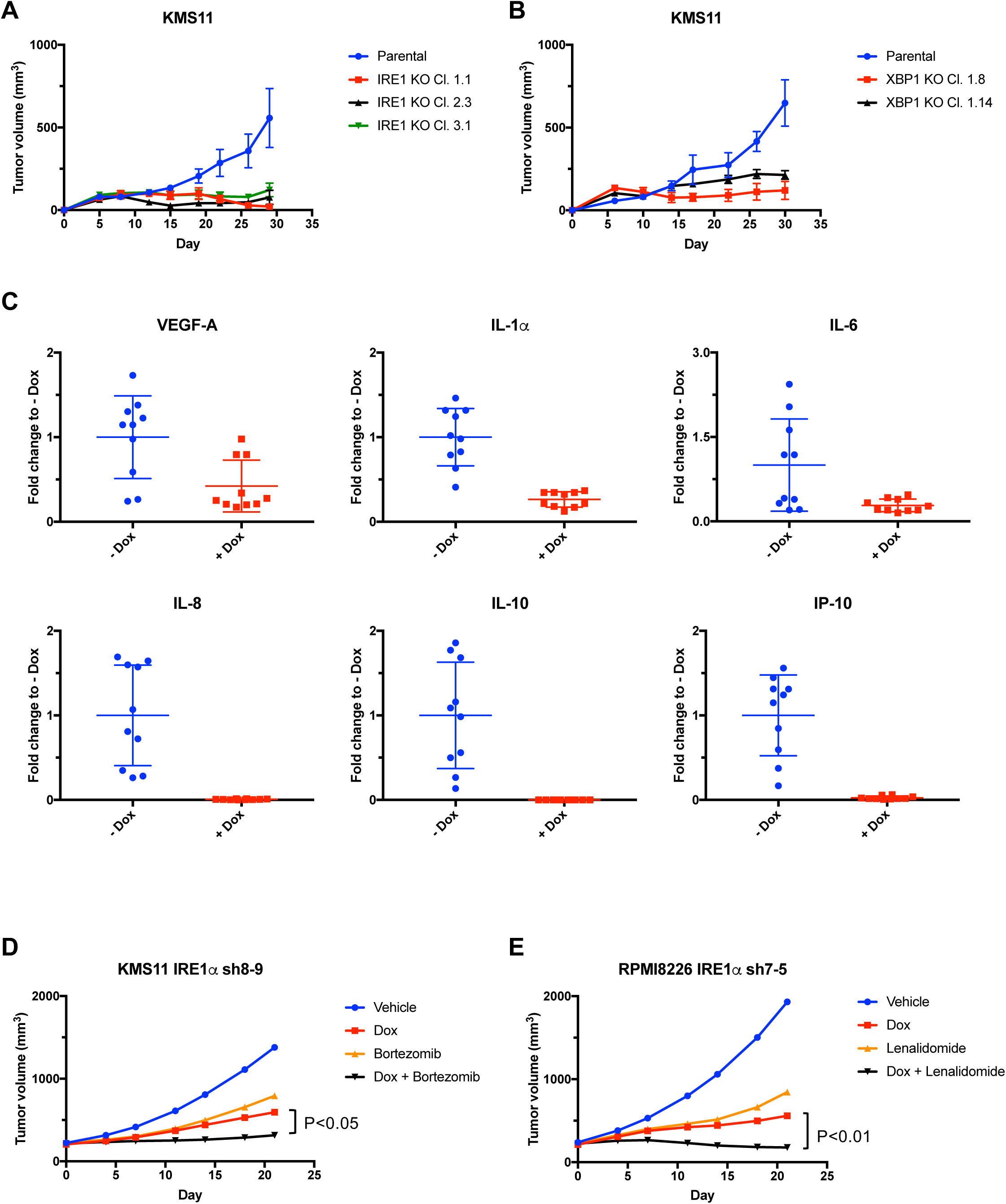
Genetic disruption of IRE1α or XBP1 attenuates growth of subcutaneous human MM xenografts in mice. (***A*** and ***B***) The IRE1α (***A***) or XBP1 (***B***) genes were disrupted by CRISPR/Cas9 technology in KMS11 cells. Parental or corresponding knockout (KO) clones were injected subcutaneously into CB.17-SCID mice and monitored for tumor growth over 29 or 30 days, respectively. (***C***) Mice bearing subcutaneous RPMI8226 tumor xenografts were treated with Dox (0.5 mg/kg) for 21 days in drinking water and analyzed for concentrations of indicated cytokines and chemokines by ELISA. (***D*** and *E*) KMS11 (***D***) or RPMI8226 (***E***) cells stably transfected with Dox-inducible shRNAs against IRE1α were inoculated subcutaneously into C.B17-SCID mice and allowed to establish tumors of ~200 mm^3^ in volume. Mice were then treated with sucrose or Dox in drinking water and tumors were monitored for 21 days. Alternatively, mice with established tumors were treated with the proteasome inhibitor bortezomib (0.75 mg/kg IV), alone or in combination with Dox (***D***); or the IMiD lenalidomide (50 mg/kg IP), alone or in combination with Dox (***E***). Individual tumor data are shown in **SI Appendix**, **Fig. S2*F*** and ***G***.

To examine the relative importance of XBP1s for MM tumor growth, we disrupted the XBP1 gene by CRISPR/Cas9 in KMS11 cells. Similar to the IRE1α KO clones, two independent XBP1 KO clones failed to grow appreciably upon subcutaneous injection into Cb.17-SCID mice, while parental WT cells formed tumors as expected (**Fig. 2*B*; SI Appendix, Fig. S2*C***). Thus, *in vivo* growth of KMS11 xenografts requires XBP1s. Mechanistically, IRE1α KO decreased the *in vitro* mRNA levels of several XBP1s target genes encoding key components of the cellular ERAD machinery (14, 17), *i.e*., the E3 ubiquitin ligase SYVN1, the E2 ubiquitin-conjugating enzyme UBE2J1, and factors required for the recognition and extraction of terminally misfolded proteins from the ER, such as EDEM1, DERL2, VIMP, DNAJC10, and ERLEC1 (**SI Appendix, Fig. S2*D***). It also attenuated secretion and increased cellular retention of IgG by cultured MM cells (**SI Appendix, Fig. S2*E***). Furthermore, *in vivo* IRE1α depletion in a subcutaneous RPMI8226 xenograft model inhibited secretion into the bloodstream of several cytokines, including vascular endothelial growth factor (VEGF), interleukin (IL)-6, IL-10 and IL-1α, as well as of certain chemokines, including IL-8 (CXCL8) and interferon-inducible protein (IP)-10 (CXCL10) (**Fig. 2*C***). The perturbation of both ERAD and of secretory functions in MM cells lacking IRE1α may compromise their *in vivo* growth (23, 41).

To ascertain whether IRE1α depletion alone or in combination with standard anti-MM therapies affects growth of pre-established tumors, we allowed subcutaneously implanted KMS11 or RPMI8226 cells carrying Dox-inducible IRE1α shRNAs to form palpable tumors of ~200 mm^3^, and only then induced depletion by Dox treatment. IRE1α knockdown substantially suppressed tumor progression, in conjunction with a marked decrease in XBP1s protein levels; this led to 61% tumor-growth inhibition (TGI) in KMS11 and 70% TGI in RPMI8226 xenografts (**Fig. *2D* and *E*; SI Appendix, Fig. S2*F-I*)**. Furthermore, treatment in the KMS11 model with the maximum tolerated dose (MTD) of the PI bortezomib led to 54% TGI, while the combination of IRE1α knockdown with bortezomib afforded 91% TGI (p<0.05 compared to IRE1α knockdown alone) (**Fig. 2*D*; SI Appendix, Fig. S2*F***), indicating strong tumor attenuation. Similarly, treatment in the RPMI8226 model with the MTD of the IMiD lenalidomide led to 61% TGI, while the combination of IRE1α depletion with lenalidomide achieved 110% TGI (p<0.01 compared to IRE1α knockdown alone) (**Fig. 2*E*; SI Appendix, Fig. S2*G***), indicating complete tumor suppression or regression. Together, these results show that genetic disruption of IRE1α markedly inhibits MM tumor initiation and progression, and increases sensitivity to established agents. Clinically, the perturbation of IRE1α has the potential to cooperate significantly with existing treatment modalities to enhance the efficacy of anti-MM therapy.

### Small-molecule inhibition of IRE1α kinase attenuates subcutaneous and orthometastatic growth of human MM xenografts

Next, we investigated whether pharmacologic inhibition of IRE1α could recapitulate the impact of genetic disruption on MM tumor growth. Because XBP1s depletion through direct IRE1α RNase inhibition can lead to hyper-phosphorylation of the kinase domain (20, 42), we chose to block IRE1α further upstream, at its kinase level. To test whether IRE1α auto-phosphorylation controls RNase activation in MM cells, we transfected KMS11 IRE1α KO cells with cDNA expression plasmids encoding WT or mutant variants of IRE1α enzymatically deficient in kinase activity (D688N) or auto-phosphorylation on the kinase-activation loop (S724A S726A S729A triple mutant). Upon ER stress, cells expressing WT IRE1α, but not the kinase-dead or auto-phosphorylation-deficient mutants, displayed elevated production of XBP1s mRNA and protein (**Fig. 3*A***). Thus, disruption of either the kinase function or the auto-phosphorylation sites of IRE1α in MM cells blocks RNase activation and XBP1s production.

**Fig. 3.**
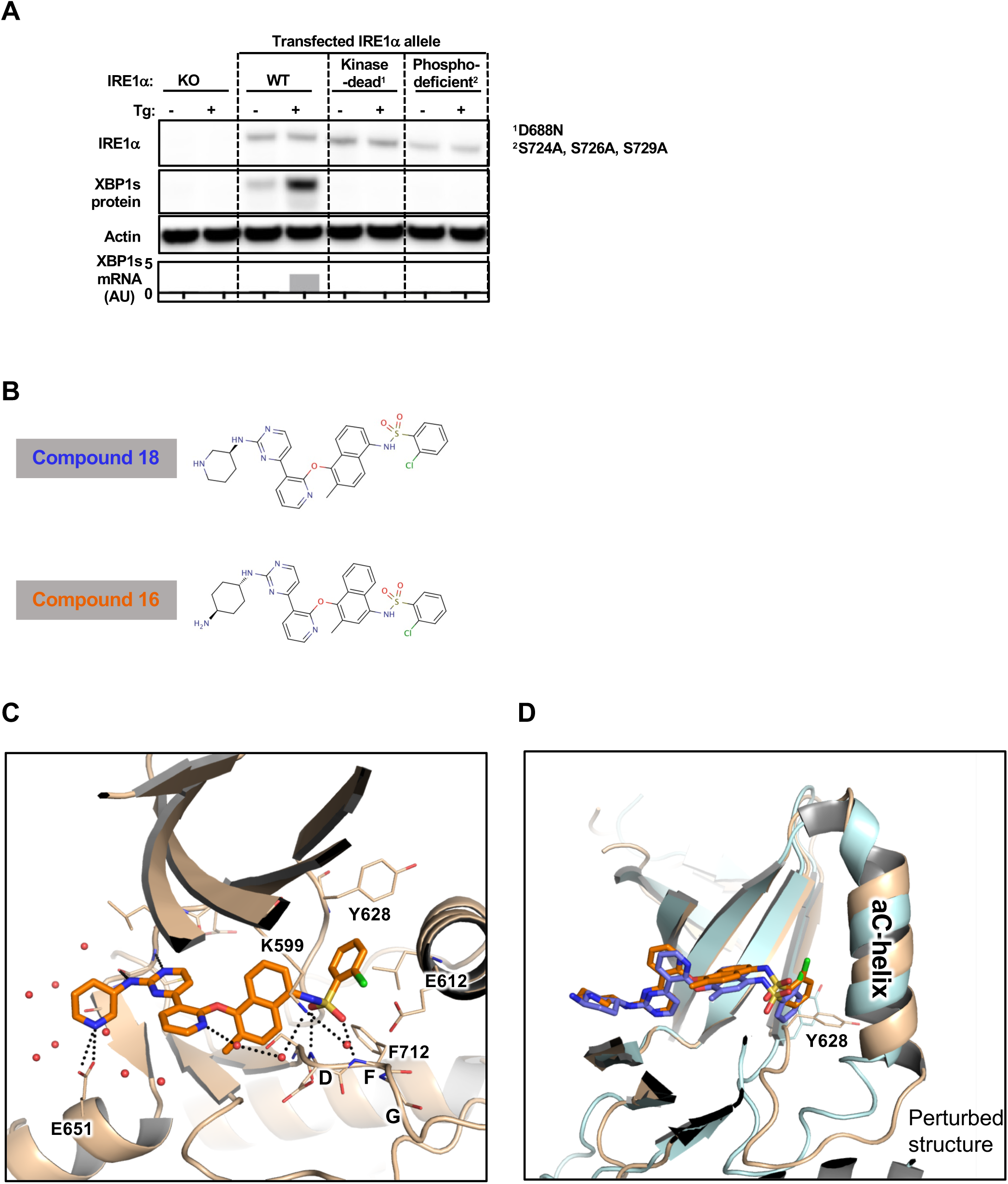
Importance of IRE1α kinase in RNase activation and co-crystal structure of IRE1α with compound 18 as compared to 16. (***A***) KMS11 IRE1α KO cells were stably transfected with expression plasmids encoding: Wildtype (WT) IRE1α, or a “kinase-dead” D688N mutant of IRE1α, or an “auto-phosphorylation-deficient” S724A, S726A, S729A triple mutant of IRE1α. Cells were then incubated in the absence or presence of Tg (100 nM) for 3 hours, and analyzed either by IB for levels of the indicated proteins, or by RT-QPCR for mRNA levels of XBP1s. (***B***) Chemical structure of **18** and **16** (36). (***C***) A close-up view of the crystal structure of compound **18** in complex with the kinase-RNase portion of IRE1α. The protein is rendered in ribbons, with key residues in the ligand-binding pocket shown as sticks. Water molecules near the ligand are shown in red spheres. Black dash lines indicate hydrogen bonding interactions. (***D***) Comparison between the co-crystal structures of compound **18** (colored in wheat and orange versus **16** (PDB entry: 4U6R, colored cyan and blue) bound to IRE1α. The C-terminal end of the αC-helix displays significant conformational changes between the two structures.

Harrington et al (36) identified kinome-selective inhibitors of IRE1α kinase, including compounds **16** and **18**. We synthesized both molecules and confirmed their binding to a recombinant IRE1α protein comprising the kinase and RNase domains, and their ability to inhibit its RNase activity toward a synthetic XBP1-based RNA substrate, as well as cellular IRE1α activity measured by an XBP1s-luciferase reporter assay (**SI Appendix, Fig. S3*A***) (11, 24). We compared the kinase selectivity of these compounds by testing 220 kinases via KinomeScan™. Compound **18** displayed significantly better selectivity than **16**, with > 70% inhibition of only one off-target kinase (JNK2), as compared to eight for **16** (**SI Appendix, Fig. S3*B***). Of note, two different JNK-specific inhibitors, SP600125 and JNK-IN-8, did not impact growth of KMS11 cells at concentrations of up to 10 μM (data not shown). Furthermore, mRNA expression of JNK2 in RPMI8226 and OPM2 cells was relatively low as compared to most other cell lines in the CCLE dataset (**SI Appendix, Fig. S3*C***), suggesting that any off-target inhibition of JNK2 by **18** in these MM cell lines is unlikely to be functionally significant. Quantitative PCR analysis further demonstrated that **18** inhibited the constitutive IRE1α-mediated XBP1s production observed in RPMI8226 cells as well as ER-stress-induced XBP1s mRNA generation and RIDD activity toward DGAT2 mRNA in KMS11 cells, with half-maximal inhibitory concentrations (IC_50_) of 30-60 nM (**SI Appendix, Fig. S3*D-G*)**.

To gain structural insight into the interaction of compound **18** with its target, we co-crystallized it with the purified recombinant IRE1α kinase-RNase protein and determined an X-ray structure at 2.20 Å resolution. Compound **18** binds in the ATP docking site (**Fig. 3*C*; SI Appendix, Fig. S3*G***), consistent with its ability to act as a kinase inhibitor of IRE1α. The aminopyrimidine anchors at the hinge and delivers the chloro-phenyl tail moiety to the kinase back-pocket. The sulfonamide forms hydrogen bonds with the Asp, Phe, Gly (DFG) backbone in a DFG-in conformation and accepts a hydrogen bond from the catalytic Lys residue, K599. The Lys-Glu salt-bridge typically seen in the active state of kinases is absent in this structure, as K599 and E612 are separated by 5.4 Å. The combined effects of back-pocket binding and salt-bridge disruption may induce critical structural changes throughout the cytoplasmic region that ultimately afford allosteric inhibition of the RNase. This ligand-binding mode is reminiscent of the interaction of **16** with IRE1α (PDB entry 4U6R) (36). However, the 1,4 substituted naphthyl linker of **18** pulls back from the kinase N-lobe by ~1.0 Å as compared to the 1,5 substituted naphthyl linker of **16**. Further comparison reveals that **18** displaces the C-terminal end of the Cα-helix to a greater extent than does **16** (**Fig. 3*D***), where residue Y628 shows the most difference in sidechain conformation. Although we cannot rule out that crystal packing may influence this, structural changes in the Cα-helix may contribute to the improved selectivity of **18** against IRE1α: We therefore chose the latter molecule as a tool for further studies.

Compound **18** was not suitable for oral dosing (data not shown). However, upon intraperitoneal dosing at 30 mg/kg once or twice per day in C.B-17 SCID mice, **18** achieved initial plasma concentrations of ~5 μM, and remained above 0.1 μM for ~8 hr (**SI Appendix, Fig. S4*A***). These data suggested potentially sufficient exposure to this compound to attain significant, though perhaps incomplete, IRE1α inhibition *in vivo*. Comparable to the effect of IRE1α shRNA depletion, treatment of mice bearing pre-established KMS11 or OPM2 tumors with **18** over 23 or 11 days, respectively, led to a substantial reduction in XBP1s protein, in conjunction with 51% or 70% TGI, respectively (**Fig. 4*A* and B; SI Appendix, Fig. S4*B-E***). Thus, pharmacologic IRE1α kinase inhibition recapitulated the impact of shRNA-based IRE1α disruption on growth of MM xenografts.

**Fig. 4.**
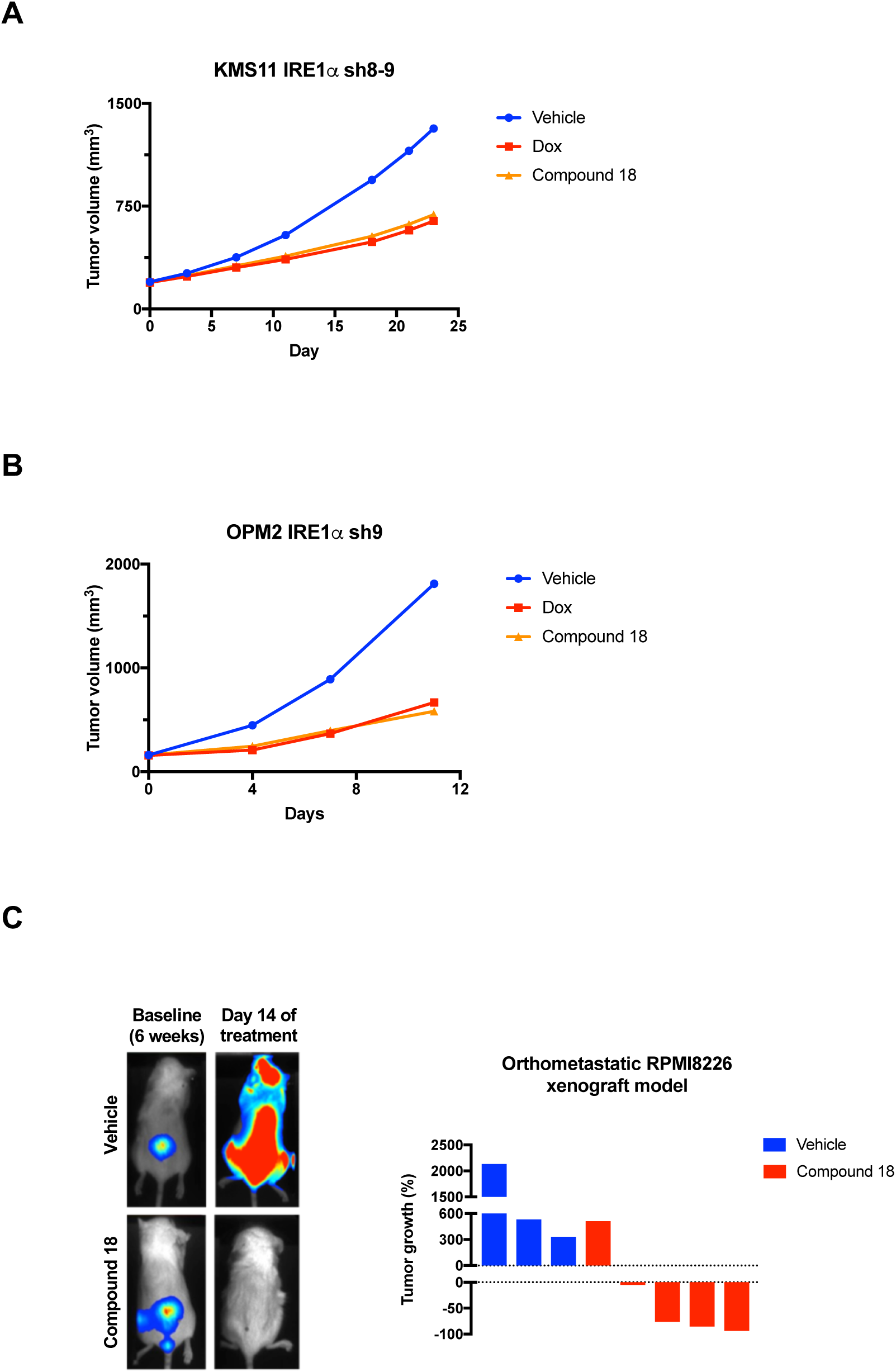
Small-molecule inhibition of IRE1α kinase attenuates subcutaneous and orthometastatic growth of human MM xenografts in mice. (**A**) KMS11 cells stably transfected with plasmids encoding Dox-inducible shRNAs against IRE1α were inoculated subcutaneously into C.B17-SCID mice and allowed to establish tumors of ~200 mm^3^. Mice were then treated with either vehicle, Dox in the drinking water (0.5 mg/kg), or compound **18** (30 mg/kg) intraperitoneally twice per day and monitored for tumor growth over 24 days. Individual tumor data are shown in **SI Appendix, Fig. S4*C***. (B) OPM2 cells stably transfected with plasmids encoding Dox-inducible shRNAs against IRE1α were inoculated subcutaneously into C.B17-SCID mice and allowed to establish tumors of ~160 mm^3^. Mice were then treated as in ***A*** with either vehicle, Dox in the drinking water, or compound **18** intraperitoneally once per day and monitored for tumor growth over 11 days. Individual tumor data are shown in **SI Appendix, Fig. S4*E***. (***C***) RPMI8226 cells expressing plasmids encoding mCherry and luciferase were injected intravenously via the tail vein of NSG mice and tumors were allowed to establish in the bone marrow over a period of 6 weeks. Tumor burden was monitored by in-life imaging of luminescence. After 6 weeks, mice were grouped out based on similar tumor burden, treated with vehicle (n=3) or compound **18** (30 mg/kg, intraperitoneally, twice per day, n=5) for 2 weeks, and analyzed again for tumor burden. One control mouse died during anesthesia and one treated mouse was euthanized due to weight loss. Luminescence images of representative mice are depicted on the left. The tumor burden of each mouse is depicted as % tumor growth on day 14 (at the end of 8 weeks) as compared to day 0 of treatment (at the end of 6 weeks).

We then turned to a more stringent orthometastatic model of MM, in which luciferase and mCherry double-labeled RPMI8226 cells, injected into the tail vein of NSG mice, develop widespread malignant disease with bone marrow involvement over a period of 6 weeks (SI Appendix, Fig. *S4F*) (43). Daily treatment of mice bearing established malignant disease with compound 18 over two subsequent weeks led to a substantial reduction in tumor burden, as evident by diminished luminescence (**Fig. 4*C***): Whereas 3/3 control mice displayed tumor progression over baseline, only 1/5 **18**-treated mice showed tumor progression, while another 1/5 exhibited tumor stasis, and 3/5 showed substantial tumor regression. Thus, pharmacologic inhibition of IRE1α kinase *in vivo* can disrupt growth of MM xenografts not only in the subcutaneous setting but also in the more clinically relevant orthometastatic bone marrow microenvironment.

### IRE1α kinase inhibition reduces viability of patient-derived MM cells while sparing normal cells

MM cell lines may acquire further genetic or epigenetic alterations upon prolonged passage that could diverge from primary MM cells. Therefore, to gain a more direct appraisal of the importance of IRE1α for primary MM cell survival, we tested the effect of compound **18** on viability of primary CD138^+^ MM cells, from bone marrow or peripheral blood samples donated by MM patients treated in the USA and EU (**SI Appendix, Fig. S5*A***). Incubation over two days with **18** led to marked reductions in viability of the malignant CD138^+^ MM cells, but not the associated non-malignant CD138^−^ cells, in the majority of MM samples (**Fig. 5*A* and *B***). Samples in both cohorts from newly diagnosed patients as well as subjects whose MM relapsed after 1-4 prior lines of therapy showed dose-dependent sensitivity to **18** (**Fig. 5*C* and *D***). Four additional relapsed MM samples did not show significant loss of viability (data not shown). Importantly, exposure to **18** did not reduce the viability of CD138^+^ cells from three nonmalignant bone marrow aspirates (**Fig. 5*E***). Thus, IRE1α kinase inhibition can selectively disrupt survival of primary malignant MM cells while sparing nonmalignant marrow cells, including plasma cells. The impact on both naiive and post-treatment relapsed MM samples suggests that IRE1α inhibition has the potential to provide clinical benefit across multiple lines of therapy.

**Fig. 5.**
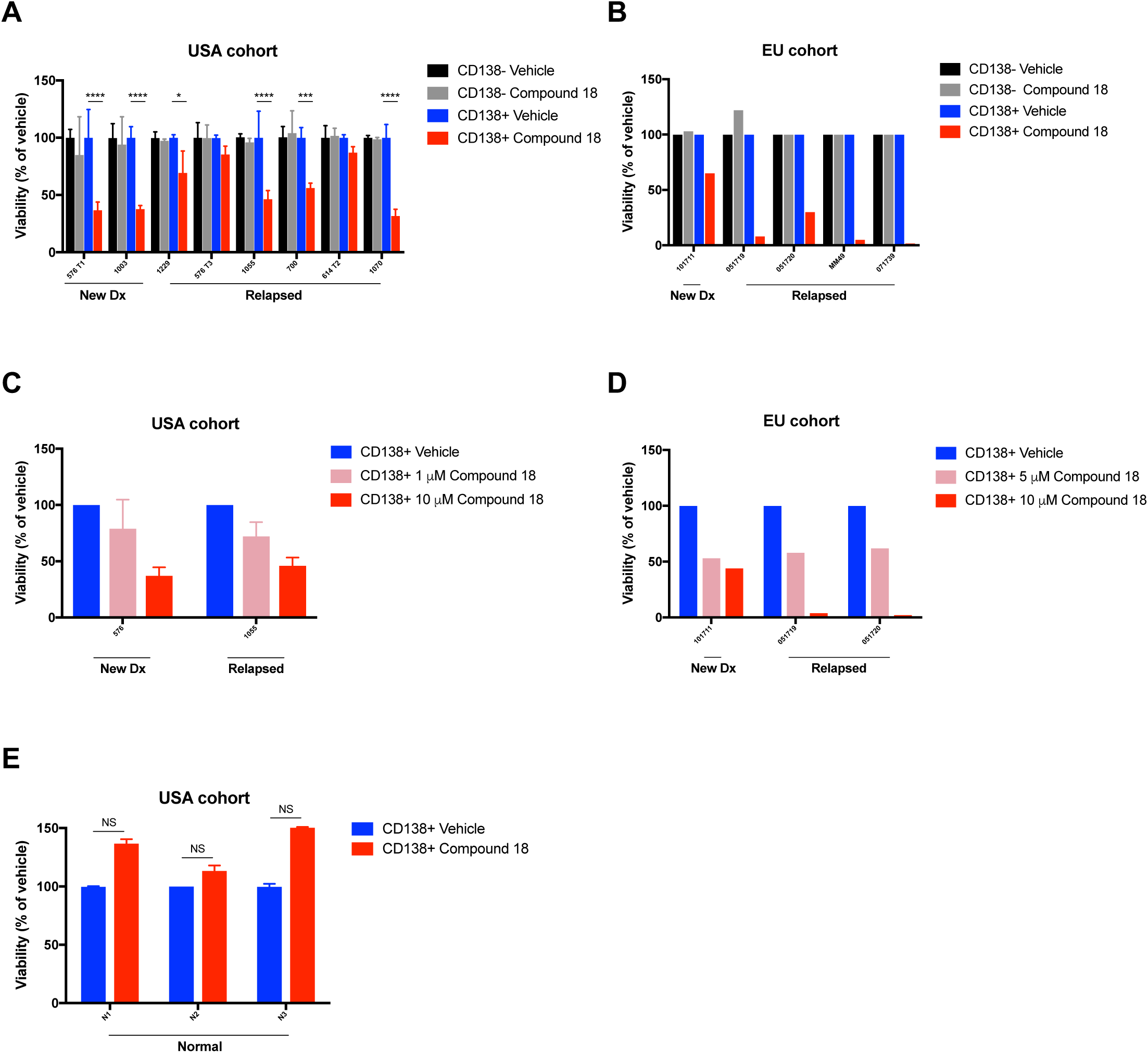
Small-molecule inhibition of IRE1α kinase reduces viability of CD138^+^ MM cells in patient-derived samples without disrupting CD138^−^ cells. Bone marrow aspirates or peripheral blood obtained in the USA (***A, C***) or EU (***B, D***) from patients with newly diagnosed or relapsed MM were cultured for 48 hr (***A-C***) or 72 hr (***D***) in the absence or presence of compound **18** at the indicated concentration. Samples were then analyzed for viability by flow cytometry, with gating on CD138^+^ or CD138^−^ cells. Nonmalignant bone marrow aspirates (n=3) were similarly tested and are depicted for comparison (***E***). Data represent mean ± SD of triplicate determinations except where triplicates were not possible due to insufficient sample size. Statistical analysis in panels **A** and **B** was performed by ANOVA. NS, not significant. Further information about age, gender, disease state, cytogenetics, and prior treatments is included in **SI Appendix, Fig. S5*A***.

Inducible gene-knockout studies in mice have suggested that the IRE1α-XBP1s pathway may support insulin secretion by pancreatic cells (44, 45). To examine whether pharmacologic IRE1α kinase inhibition would affect these cell types, we first verified the ability of compound **18** to inhibit XBP1s induction in human pancreatic islet 3D microtissues, which contain all the endocrine cell types and can retain viability and function in culture for up to 4 weeks (46). At 2.4 μM, **18** suppressed Tg-induced XBP1s production to baseline levels (**Fig. 6*A***), confirming effective IRE1α pathway inhibition. Importantly, **18** did not decrease viability, nor did it perturb glucose-stimulated insulin secretion even at higher concentrations up to 7.5 μM (**Fig. 6*B* and *C***). We obtained similar results with rat pancreatic microislets (**SI Appendix**, **Fig. S6*A*** and ***B***). Together, these results suggest that IRE1α kinase inhibition may achieve significant MM tumor disruption without overt negative effects on pancreatic endocrine homeostasis.

**Fig. 6.**
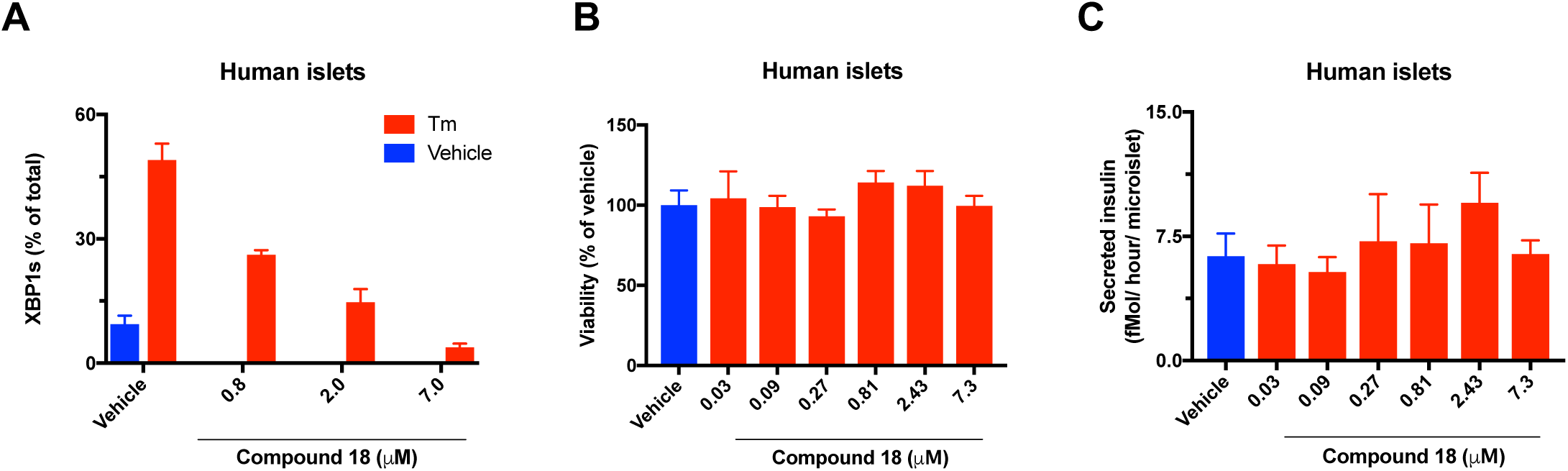
IRE1α kinase inhibition preserves survival and insulin secretion by pancreatic islet 3D microtissues. (***A-C***) Human pancreatic islets were isolated, dissociated into single cells, replated in microtiter wells (1000 cells/drop), and allowed to form 3D microtissues of ~120 μΜ in diameter over 7 days using inSphero™ technology. Microtissues (n=5 per treatment) were then (***A***) treated for 24 hr with tunicamycin (5 μg/ml) in the absence or presence of compound **18** at the indicated concentrations, lysed, and then analyzed for XBP1s/(XBP1s+XBP1u) mRNA levels by RT-QPCR; or (***B*** and ***C***) incubated for 7 days in the absence or presence of **18** at the indicated concentrations, and then (**B**) analyzed for cell viability by CellTiterGlo™ assay; or (***C***) challenged with glucose (16.7 mM) for 1 hr and analyzed for insulin secretion by ELISA.

## Discussion

It has been proposed that MM cells may coopt the IRE1α-XBP1s pathway to mitigate persistent ER stress, caused by excessive Ig production and a nutrient/oxygen-poor bone marrow microenvironment (27). However, recent studies have raised significant doubt concerning the validity of IRE1α as a potential therapeutic target: Lowered levels of XBP1s correlated with PI resistance in MM cells (35), and IRE1α kinase inhibition blocked XBP1s yet did not affect MM cell viability in 2D culture (35, 36). Moreover, loss-of-function studies directly and specifically addressing the importance of IRE1α’s kinase module for MM growth have been lacking. Although work based on salicylaldehyde small-molecule RNase inhibitors supported a protumorigenic role of IRE1α in MM (31, 32), a key caveat is that such compounds are highly protein-reactive, and their selectivity *versus* other RNases and other targets is likely to be poor (37).

To interrogate the requirement of IRE1α for MM growth, we employed a series of strategies to disrupt it at the gene, transcript, or kinase level, in diverse model systems. Our *in vitro* studies showed that IRE1α depletion by shRNA markedly attenuates growth of several MM cell lines in 3D spheroid settings—a scenario that was not previously investigated. Both IRE1α KO and XBP1s KO by CRISPR/Cas9 in KMS11 MM cells profoundly disrupted their ability to form subcutaneous tumor xenografts in mice. These findings demonstrate a critical requirement for the IRE1α pathway for *in vivo* MM growth, while other XBP1s-independent functions of IRE1α such as RIDD and JNK activation may be less crucial in this context. IRE1α depletion by shRNA also inhibited growth of pre-formed subcutaneous KMS11 and RPMI8226 tumor xenografts, implicating IRE1α in promoting not only tumor initiation but also tumor progression. Mechanistically, IRE1α knockdown decreased mRNA expression of specific ERAD components known to be induced by XBP1s, and it compromised the ability of MM cells to secrete Ig as well as several important cytokines and chemokines, some of which have previously been shown to support malignant plasma cell growth *in vitro* and *in* vivo (1, 23, 33, 41). Remarkably, the extent of TGI was comparable between IRE1α knockdown and the well-established frontline MM therapy agents bortezomib or lenalidomide. Furthermore, combination of IRE1α depletion with bortezomib or lenalidomide significantly increased the extent of TGI as compared to respective monotherapies. Together, these results suggest that IRE1α inhibition may provide clinical benefit, either alone or in combination with other MM therapies known to disrupt protein homeostasis (33, 47).

To examine more directly the requirement for IRE1α’s kinase moiety, we first confirmed its importance for RNase activation by mutational perturbation of the kinase catalytic core or its target autophosphorylation sites. We then evaluated two compounds that bind to IRE1α’s ATP docking site and exert allosteric inhibition of RNase activation (36). One of these showed an improved ability to displace the Cα helix in the kinase domain, with excellent selectivity toward IRE1α versus 220 other kinases. This compound displayed sufficient tolerability and plasma exposure upon intraperitoneal administration *in vivo* to enable substantial inhibition of XBP1s production in tumors. Moreover, it significantly attenuated subcutaneous growth of KMS11 and OPM2 xenografts. Thus, pharmacologic inhibition of IRE1α via its kinase moiety recapitulated the impact of genetic IRE1α disruption on MM tumor growth.

To address the importance of IRE1α for MM growth in a more clinically relevant microenvironment, we used an orthometastatic model, in which malignant MM cells injected intravenously home to the bone marrow to disseminate malignant disease. Treatment with the IRE1α kinase inhibitor in this setting led to tumor stasis or regression in most of the compound-treated animals, as compared to aggressive tumor progression in vehicle-treated controls. Thus, MM cells require IRE1α activity to sustain advanced malignant growth based in the bone marrow microenvironment.

Establishing the excellent kinase selectivity of compound **18** afforded us a unique opportunity to examine more reliably the impact of specific IRE1α inhibition on patient-derived MM cells. Remarkably, the compound caused a substantial reduction in viability of malignant CD138^+^ cells in most of the MM patient samples. Importantly, these included newly diagnosed tumors as well as some tumors that relapsed after up to four lines of prior therapy with PI and/or IMiD and/or other agents. In contrast, **18** did not significantly reduce viability of accompanying nonmalignant CD138^—^ cells in the same MM aspirates; it also spared both CD138^+^ plasma cells and CD138^—^ cells in nonmalignant bone marrow samples. Furthermore, while achieving complete XBP1s suppression, 18 disrupted neither the overall viability of human and rat pancreatic microislets, nor their capacity to secrete insulin in response to glucose challenge. Taken together, these results suggest that malignant MM cells may harbor a greater dependency on the IRE1α-XBP1s pathway than do nonmalignant secretory cell types including normal bone marrow plasma cells, highlighting the importance of this pathway as a unique vulnerability that could be clinically exploited to treat MM across multiple stages. An effort to develop orally available compounds to target IRE1α kinase and a more comprehensive safety analysis in suitable model organisms would be needed in order to enable future human testing of IRE1α inhibitors in clinical trials.

In conclusion, our studies provide definitive preclinical evidence validating IRE1α as a potential therapeutic target for MM. IRE1α may play an important role in augmenting malignant growth of MM cells by enabling their adaptation to chronic ER stress through elevated ERAD capacity. IRE1α may also support the secretion of cytokines and chemokines that enable survival and growth of malignant MM cells in their metabolically restrictive bone marrow microenvironment. Finally, our findings provide proof of concept that the kinase domain of IRE1α is an effective and potentially safe lever for small-molecule inhibition of this dual-function enzyme both *in vitro* and *in vivo*. This work therefore presents a compelling rationale to develop clinical-grade IRE1α kinase-disrupting agents for MM therapy.

## Material and Methods

Detailed methods are provided in SI Appendix.

### Cell culture and experimental reagents

KMS11, RPMI8226, OPM2, H929, KMS27, MOLP8, LP1, U266.B1, UTMC2, KMM1, KMS28PE and MOLP2 cells were obtained from ATCC, JCRB, or DSMZ, authenticated by short tandem repeat (STR) profiles, and tested to ensure mycoplasma free within 3 months of use. All cell lines were cultured in RPMI1640 media supplemented with 10% (v/v) fetal bovine serum (FBS, Sigma), 2 mM glutaMAX (Gibco) and 100 U/ml penicillin plus 100 μg/ml streptomycin (Gibco).

Thapsigargin (Sigma) was used at a concentration of 100 nM, tunicamycin (Sigma) at 5 μg/ml. Compound 16/ **16** and compound 18/ **18** (36) were dissolved in DMSO for cellular experiments, and used at the indicated concentrations. Antibodies (Abs) for IRE1α (#3294), β-actin (#3700) and GAPDH (#5174) were from Cell Signaling Technology. Ab for human IgG (#109489) was from Abcam. Abs for XBP1s and pIRE1 (21) were generated at Genentech. Secondary antibody (#711-035-152) was from Jackson Laboratories.

### 3D spheroid proliferation assays

RPMI8226 IRE1α sh7-5-mCherry, OPM2 IRE1α sh9-mCherry and KMS11α IRE1 sh8-9-mCherry cells were pre-treated with 0.5 μg/ml doxycycline for 3 days before plating 1000 cells/ well in ultra-low attachment (ULA) 96-well plates (Corning). Single tumor spheroids were formed by centrifugation (1,000 rpm) for 10 min according to manufacturer’s protocol. Spheroids were imaged using a real-time imaging system (IncuCyte™, Essen Bioscience, Ann Arbor, Michigan). Frames were captured at 4-hr intervals using a 4x or 10x objective and red fluorescence was detected.

For Matrigel assays, KMS11 IRE1α sh8-9 cells were pre-treated with 0.5 μg/ml doxycycline for 3 days before plating 1000-5000 cells/well on 50 μl/well of Matrigel (Corning) into 96 well plates according to manufacturer’s protocol. Spheroids images were captured at 4-hr intervals using a 10x objective. Cell viability was assessed using an ATP-consumption assay (CellTiter-Glo^®^ 3D, Promega) and measured in a luminescence reader (Envision, Perkin Elmer). Cultures were maintained at 37 °C throughout and run at least in triplicates. Values well were pooled and averaged across all replicates.

### Pancreatic islet 3D microtissue assays

Human and rodent 3D InSight™ pancreatic islet microtissues (InSphero AG) were produced from reconstituted dispersed human or rat pancreatic islet cells in a modified manner as described previously (46) retaining the composition of α, β and δ cells representative of normal endocrine pancreatic islets. Cells were plated in microtiter wells (1000 cells/drop), and allowed to form 3D microtissues of ~120 μM in diameter over 7 days (n=5 per treatment). Microtissues were incubated for 7 days with serial dilutions of compound **18** or vehicle control, and then viability analyzed by CellTiter-Glo^®^ (Promega), or insulin secretion analyzed after glucose challenge for 2 hours (16.7 μM) by ELISA.

### Human MM samples

The effect of compound **18** on viability of MM or normal cells was measured after a 48 hr treatment in *ex vivo* culture of bone marrow aspirates or blood samples from MM patients or from normal bone marrow donors. For cell death assays, mononuclear cells (MNC) obtained after separation on Ficoll density gradient where cultured in RPMI1640 media supplemented with 5% FCS and 3 ng/ml IL-6, with the indicated concentrations of compound **18** for 48 hr. MM cells were then identified using CD138-PE staining and cell death was assessed by the loss of CD138 staining as previously described (48). MM or normal plasma cells were identified as CD19^−^, CD45^−^/dim, CD38^+^, CD138^+^, and CD46^+^.

### Subcutaneous xenograft growth and efficacy studies

All procedures were approved by and conformed to the guidelines and principles set by the Institutional Animal Care and Use Committee (IACUC) of Genentech and were carried out in an Association for the Assessment and Accreditation of Laboratory Animal Care (AAALAC)-accredited facility.

For tumor growth studies, 10×10^6^ KMS11 WT, IRE1α KO or XBP1 KO clones, respectively, were suspended in HBSS, admixed with 50% Matrigel (Corning) to a final volume of 100 μl, and injected subcutaneously in the right flank of 6 to 8-week old female CB-17 SCID mice.

For efficacy studies, 10×10^6^ KMS11 IRE1α sh8-9, RPMI8226 IRE1α sh5-7 or OPM2 IRE1α sh9 cells were prepared and subcutaneously inoculated as outlined above. Tumors were monitored until they reached a mean tumor volume of approximately 150 to 300 mm^3^. For efficacy studies of IRE1 shRNA knockdown in combination with standard of care agents, bortezomib (Velcade^®^, Millennium Pharmaceuticals) or lenalidomide (Revlimid^®^, Celgene Corp.), animals were randomized into the following treatment groups: (i) 5% sucrose water (provided in drinking water, changed weekly); (ii) doxycycline (0.5 mg/ml, dissolved in 5% sucrose water, changed 3x/ week); (iii) bortezomib (0.75 mg/kg, 100 μL total, intravenously, twice per week) or lenalidomide (50mg/kg, 100 μL total, intraperitoneally, once daily for 5 consecutive days), respectively; and (iv) combination of doxycycline plus bortezomib, or doxycycline plus lenalidomide, respectively.

For the compound **18** efficacy studies, animals were randomized into one of three following treatment groups: (i) vehicle controls (35% PEG400 and 10% EtOH in water, 100 μL total, intraperitoneally, dosed once daily) and 5% sucrose water (provided as drinking water, changed weekly); (ii) doxycycline (0.5 mg/ml, dissolved in 5% sucrose water, changed 3x/ week); (iii) compound **18** (30 mg/kg, 100 μL total, intraperitoneally, dosed QD or BID as indicated in Figure legends).

### Orthometastatic xenograft efficacy studies

For the orthometastatic xenograft model, 1×10^6^ RPMI8226-mCherry-Luc cells were injected intravenously via the tail vein of non-irradiated 8-week old female NOD/SCID/IL2rγ-/-mice (NSG, Jackson Laboratories). The animals were imaged weekly under isoflurane anesthesia 5 min after intraperitoneal luciferin injection with 200 μl of 25 mg ml^−1^ D-luciferin (Invitrogen), and imaged on a Photon Imager (BioSpace Laboratory). During image acquisition, animals continued to receive anesthesia from a nose-cone delivery system, while their body temperatures were maintained on a thermostatically controlled platform. Photon counts per min per cm^2^ of observational area were calculated and compared using M3 Vision software (BioSpace Laboratory). After 6 weeks mice were grouped out into the following treatment groups: (i) vehicle control (35% PEG400 and 10% EtOH in water, 100 μL total, intraperitoneally, QD); or (ii) compound **18** (30 mg/kg, 100 μL total, intraperitoneally, QD). After 14 days, mice were euthanized by cervical dislocation and bones harvested for fluorescence imaging using a Kodak In-Vivo FX system (Carestream Health Molecular Imaging, New Haven, Connecticut) and Carestream Molecular Imaging (MI) Software. Excitation and emission wavelengths were fixed at 550nm and 600 nm, respectively. Fluorescence images were co-registered with X-ray images using the open-source software Image J (http://rsbweb.nih.gov/ij/).

### Statistics

Statistical analysis of the results was performed by unpaired, two-tailed t-test. Comparison of three or more columns was performed using a one-way ANOVA followed by Turkey’s procedure. A P value < 0.05 was considered significant. All statistical analyses were performed using GraphPad Prism 6 (GraphPad Software, Inc.). For further information regarding statistical analysis, please see section regarding xenograft studies above.

## Supporting information

## Acknowledgments

This work was supported in part by an Irvington Postdoctoral Fellowship of the Cancer Research Institute to DAA and by a Howard Hughes Collaborative Innovation Award to PW. PW is an Investigator of the Howard Hughes Medical Institute.

We thank Rena Wang and Xiangnan Du for help with cell line generation, Brian Rabinovich for mCherry lentiviral construct design, members of the Ashkenazi and Walter labs, Marc Shuman, Stephen Gould, Matthew Wright, Shiva Malek, Daniel Sutherlin, Jessica Sims, Wendy Lee, and Ira Mellman for helpful discussions.

## Conflict of interest

J.M.H., A.L.T., S.A.M., D.A.L., M.L., Y.-C.A.C., J.Q., K.T., D.K., E.S., H.A.W., W.W., K. C., S.K., M.B., W.S., M.L., J.W., J.L., T.D.B., A.H., B.H., A.G., R.W., D.L., M.-G.B., J.R. and A.A. were employees of Genentech, Inc. during performance of this work. P.W (UCSF employee) has a patent. Rights to the invention have been licensed by UCSF to Genentech.

## Author contributions

J.M.H., A.L.T., S.A.M., D.A.L., M.L. and Y.-C.A.C. designed, performed and analyzed the experiments. J.Q. generated KMS11, OPM2 and RPMI IRE1α shRNA cell lines. K.T., D.K. and E.S. helped perform in vivo studies. H.A.W. and W.W. designed, performed and analyzed crystallographic studies. K.C., S.K. and M.B. developed assays and characterized Compound **16** and **18** in vitro. W.S., M.L. and J.W. helped generate, purify and biochemically characterize recombinant IRE1α proteins. J.L. and T.D.B. performed pharmacokinetic studies. S.A.M., A.H. and B.H. designed and generated key shRNA, gRNA and DNA constructs and corresponding genetically-engineered cell lines. A.G. and R.W. helped design, conduct and analyze in vivo bioimaging studies. D.L. helped design, perform and interpret safety studies. M.-G.B. and J.R. helped coordinate chemical synthesis and contributed to experimental design and data interpretation. M.J.V., D.W.S., P.G.-B. and M.A. designed, performed and interpreted primary MM patient sample analysis. D.A.-A. and P.W. contributed to experimental design, model development and data interpretation. J.M.H. and A.A. designed the experiments, interpreted the data and co-wrote the manuscript with input from all authors.

## Supplementary Information for Disruption of IRE1α through its Kinase Domain Attenuates Multiple Myeloma

**This PDF file includes:**

Supplemental Material and Methods

Figs. S1 to S6

References for SI reference citations

## Supplemental Material and Methods

### Generation of IRE1α shRNA-expressing cells

Cells were infected with shRNAs by lentivirus using the pG-GW-pHUSH vector system. Briefly 1×10^7^ 293T cells were plated in 6-well dishes, allowed to grow for 24 hr and transfected with lentivirus plasmids using Lipofectamine 2000 (Invitrogen) following the manufacturer’s protocol. After 6 hr the media was replaced, and after another 24 hr the virus was harvested from the cells and filtered through a 0.45 mm tube-top filter. Cleared supernatants containing the viral particles and 8 mg/ml polybreen were added to the target-cell wells, spun at 1800 rpm for 45 min at room temperature, and placed back into the incubator. After 48 hr cells were subjected to selection with with puromycin containing media. After 10 passages, cells were tested for virus by Lentivirus X p24 Rapid Titer Kit (Clonetech) then sorted using FACS and RFP+ selection.

IRE1α shRNA sequences (antisense):

shRNA5:

AGCTTTTCCAAAAAACCAAGATGCTGGAGAGATTTCTCTTGATAATCTCTCCAGCATCTTGGGGG

shRNA7:

AGCTTTTCCAAAAAAAGAACAAGCTCAACTACTTTCTCTTGATAAGTAGTTGAGCTTGTTCTGGG

shRNA8:

AGCTTTTCCAAAAAAGCACGTGAATTGATAGAGATCTCTTGAATCTCTATCAATTCACGTGCGGG

shRNA9:

AGCTTTTCCAAAAAAGAGAAGATGATTGCGATGGTCTCTTGAACCATCGCAATCATCTTCTCGGG

### CRISPR/Cas9 knockout of IRE1α and XBP1 genes

Individual IRE1α- or XBP1-specific sgRNAs were designed using a standard guide scaffold and CRISPR3 (1, 2). The gRNAs were cloned into pLKO_AIO_CMV_Cas9_mCherry, enabling co-expression of each sgRNA, Cas9, and an mCherry-based selection marker following transient transfection into target cells. A tandem array of XBP1-specific sgRNAs were designed as above and cloned into pLKO_AIO_TAN_PGK_Cas9_Puro, permitting co-expression of two XBP1-targeting sgRNAs, Cas9, and a puromycin selection marker in transfected cells. sgRNA target sequences used in this study:

IRE1α gRNA1: TCAGGAAGCGTCACTGTGC IRE1α gRNA2: GAGGACAGGCTCAATCAAA IRE1α gRNA3: TTCTCCCAGATCCTAATGA XBP1 gRNA1: TTTAGGGGTCCCGTCGGCC

Transfection was with Lipofectamine 3000 (Invitrogen) according to manufacturer’s protocol. At 24 hr after transfection, cells were washed once in PBS and resuspended in PBS media containing 3% BSA Fraction V. The cell suspension was then filtered through a 35 mm membrane followed by immediate FACS sorting using the RFP^+^ selection marker. Single cell clones (n=96) were plated and grown. Clones producing colonies were tested for proper IRE1α or XBP1 disruption by immunoblot.

### RT-qPCR

RNA was extracted with RNeasy Plus kit (Qiagen). Equal amounts of RNA were reverse transcribed and amplified by RNA-to-CT kit (Applied Biosystems). The delta-delta CT values were calculated by relating each individual CT value to its internal GAPDH control, and then normalizing to the vehicle-treatment control. Taqman primers for GAPDH, XBP1u, XBP1s, and DGAT2 were from Life Technology. RNA was purified from cells using the RNeasy Plus kit (Qiagen). Equal amounts of RNA were reverse transcribed and amplified by RNA-to-CT kit (Applied Biosystems) on the ABI QuantStudio 7 Flex Real-Time PCR System. The delta-delta CT values were calculated by relating each individual CT value to its internal GAPDH control, and then normalized to the vehicle-treatment control.

Taqman primers (Life Technologies):

GAPDH primer: Hs02758991_g1 XBP1u primer: Hs02856596_m1 XBP1s primer: Hs03929085_g1 DGAT2 primer: Hs01045913_m1

### In vitro characterization of small molecule inhibitors

Small molecule potencies were assessed in three assays of IRE1α function. Compound dilutions covering a range of concentrations from 0.2 nM to 10 μM were evaluated to determine IC_50_ values. Compound binding to the IRE1α ATP site was assessed through competition with an Alexa647-labeled staurosporine probe for binding to His-tagged IRE1α (G547-L977 D688N). Probe binding was measured as TR-FRET signal upon energy transfer between the bound probe and anti-His-allophycocyanin bound to the IRE1α. To assess inhibition of RNase activity, compound was mixed with IRE1α (Q470-L977), and 5’FAMCAUGUCC**GC**AGC**G**CAUG-3’BHQ substrate was added. Substrate cleavage was monitored kinetically as an increase in fluorescence. Cellular activity was evaluated via XBP1s-luciferase reporter assay. HEK293T cells stably transfected with the reporter construct were preincubated with compound for 2 hr and then stimulated with Tg (100 nM) for 6 hr. IRE1α-mediated cleavage and splicing of the reporter construct led to expression of luciferase, which was detected by the addition of luciferin substrate.

### Immunoblot analysis

Cells were lysed or tumor tissue mechanically disrupted in 1x RIPA buffer (Millipore) containing protease and phosphatase inhibitors (Roche), cleared by centrifugation at 13,000 rpm for 10 min, and analyzed by BCA protein assay (Thermofisher Scientific). Equal protein amounts were loaded, separated by SDS-PAGE, electro-transferred to nitrocellulose membranes using the iBLOT2 system (Invitrogen), and blocked in 5% nonfat milk solution for 2 hours. Membranes were probed with the following antibodies: IRE1α, GAPDH, β-actin (Cell Signaling Technology), XBP1s, phosphorylated IRE1α (Genentech(3)). Signal was detected using appropriate horseradish peroxidase (HRP)-conjugated secondary antibodies. All primary antibodies were used at 1:000 dilution and overnight hybridization at 4°C, followed by a one-hour incubation with horseradish peroxidase (HRP)-conjugated secondary antibodies at 1:10,000 dilution.

### Luminex and ELISA analysis

Sera from tumor-bearing mice were analyzed using Luminex Premix Panel I 32-plex (Millipore). IgG and insulin in cell supernatants were quantified using human-specific IgG (Abcam) or insulin (Mercodia) Antigen capture enzyme-linked immunosorbent assay (ELISA).

### Co-crystallographic studies

The kinase-RNase (KR) domain of hIRE1α, encoding amino acids G547-L977, was expressed as an N-terminal His6-tagged fusion protein in SF9 cells with a TEV protease cleavage site from an intracellular BEVS expression vector.

Cell pellet was resuspended in lysis buffer containing 50 mM HEPES pH 8.0, 300 mM NaCl, 10% glycerol, 1mM MgCl_2_, 1:1000 benzonase, EDTA-free PI tablets (Roche), 1mM TCEP, and 5mM imidazole. Sample was homogenized, compound **18** was added to a final concentration of 10 μM, and incubated at 4°C for 30 min. Sample was lysed by sonication, ultracentrifuged at 40,000 rpm for 45 min, and the supernatant filtered through a 0.45 μ Nalgene filter.

Cleared supernatant was batch bound to Ni-NTA Superflow beads (Qiagen) for 1 hr at 4°C with nutation. Beads were washed in lysis buffer supplemented with 25 mM imidazole, followed by protein elution in lysis buffer containing 300 mM imidazole. The eluate was concentrated and loaded onto a Hiload 16/600 Superdex 75 SEC column (GE Healthcare) equilibrated in 25 mM HEPES pH 7.8, 250 mM NaCl, 10% glycerol, 1 mM TCEP (SEC buffer). The monomeric peak was pooled and treated with lambda phosphatase for 1 hr at 37°C. Dephosphorylation to 0P was confirmed by mass spectrometry. Sample was incubated with TEV protease overnight at 4°C for tag removal. Untagged protein was isolated by passage over a Ni-NTA Superflow gravity column, followed by a wash with SEC buffer supplemented with 40mM imidazole. Flowthrough and wash fractions were pooled, diluted 1:10 in Q buffer (25 mM HEPES pH 7.8 and 1 mM TCEP), and loaded onto a 5 ml prepacked QHP column (GE Healthcare). Protein was eluted over a 50 CV gradient in Q buffers supplemented with 25 mM and 500 mM NaCl. The hIRE1α_KR + **18** complex peak was isolated and concentrated to 9.4 mg/ml for crystallography.

Crystals were generated in hanging drops with mother liquor containing 0.1 M trisodium citrate pH 5.6, 10% isopropanol, 10% PEG4000, and cesium chloride additive at 4°C. Crystals were cryoprotected in the crystallography buffer supplemented with 25% glycerol and flash frozen in liquid nitrogen. Data collection was done at ALS 5.0.2. Structure was solved to 2.2 Å.

### In vivo pharmacokinetic analysis

Pharmacokinetic properties of compound **18** were determined in 5 to 6 weeks old female CB-17 SCID mice (Charles River Laboratories) (3 per time-point for once daily (QD) or twice daily (BID) injection). The mice were administered **18** at 30 mg/kg (formulated in 35% PEG400 and 10% EtOH in water) by intraperitoneal injection QD or BID. Food and water were available *ad libitum* to all animals. Serial blood samples (15 μL) were collected by tail nick at 0.25, 0.5, 1, 2, 4, 6, or 8 hr after the administration. Blood samples were diluted with 60 μL water containing 1.7 mg/mL EDTA and kept at −80°C until analysis. Plasma concentrations of **18** were determined by a non-GLP LC/MS-MS assay.

### In vivo studies

Tumor size and body weight were measured twice per week. Subcutaneous tumor volumes were measured in two dimensions (length and width) using Ultra Cal-IV calipers (model 54 − 10 − 111; Fred V. Fowler Co.) and analyzed using Excel, version 11.2 (Microsoft), or Prism 6 (GraphPad Software, Inc.). The tumor volume was calculated with the following formula: tumor size (mm^3^) = (longer measurement × shorter measurement^2^) × 0.5. Animal body weights were measured using an Adventurer Pro AV812 scale (Ohaus Corporation). Percent weight change was calculated using the following formula: group percent weight change = [(new weight − initial weight)/initial weight] × 100. To analyze the repeated measurement of tumor volumes from the same animals over time, a mixed modeling approach was used.(4) This approach addresses both repeated measurements and modest dropouts before the end of study. Cubic regression splines were used to fit a nonlinear profile to the time courses of log2 tumor volume in each group. Fitting was done via a linear mixed-effects model, using the package “nlme” (version 3.1-108) in R version 2.15.2 (R Development Core Team 2008; R Foundation for Statistical Computing; Vienna, Austria). Tumor growth inhibition (TGI) as a percentage of vehicle was calculated as the percentage of the area under the fitted tumor volume–time curve (AUC) per day for each treatment group in relation to the vehicle control using the following formula: %TGI = %TGI = (1-[(AUC/Day)_Treatment_ ÷ (AUC/Day)_Vehicle_]) X 100. When mice reached endpoint criteria (see below) or on the last treatment day, mice were euthanized by cervical dislocation and subcutaneous xenografts harvested for immunoblot analysis.

Animals in all studies were humanely euthanized according to the following criteria: clinical signs of persistent distress or pain, significant body-weight loss (>20%), tumor size exceeding 2500 mm^3^, or when tumors ulcerated. Maximum tumor size permitted by the Institutional Animal Care and Use Committee (IACUC) is 3000 mm^3^ and in none of the experiments was this limit exceeded.

**Fig S1.**
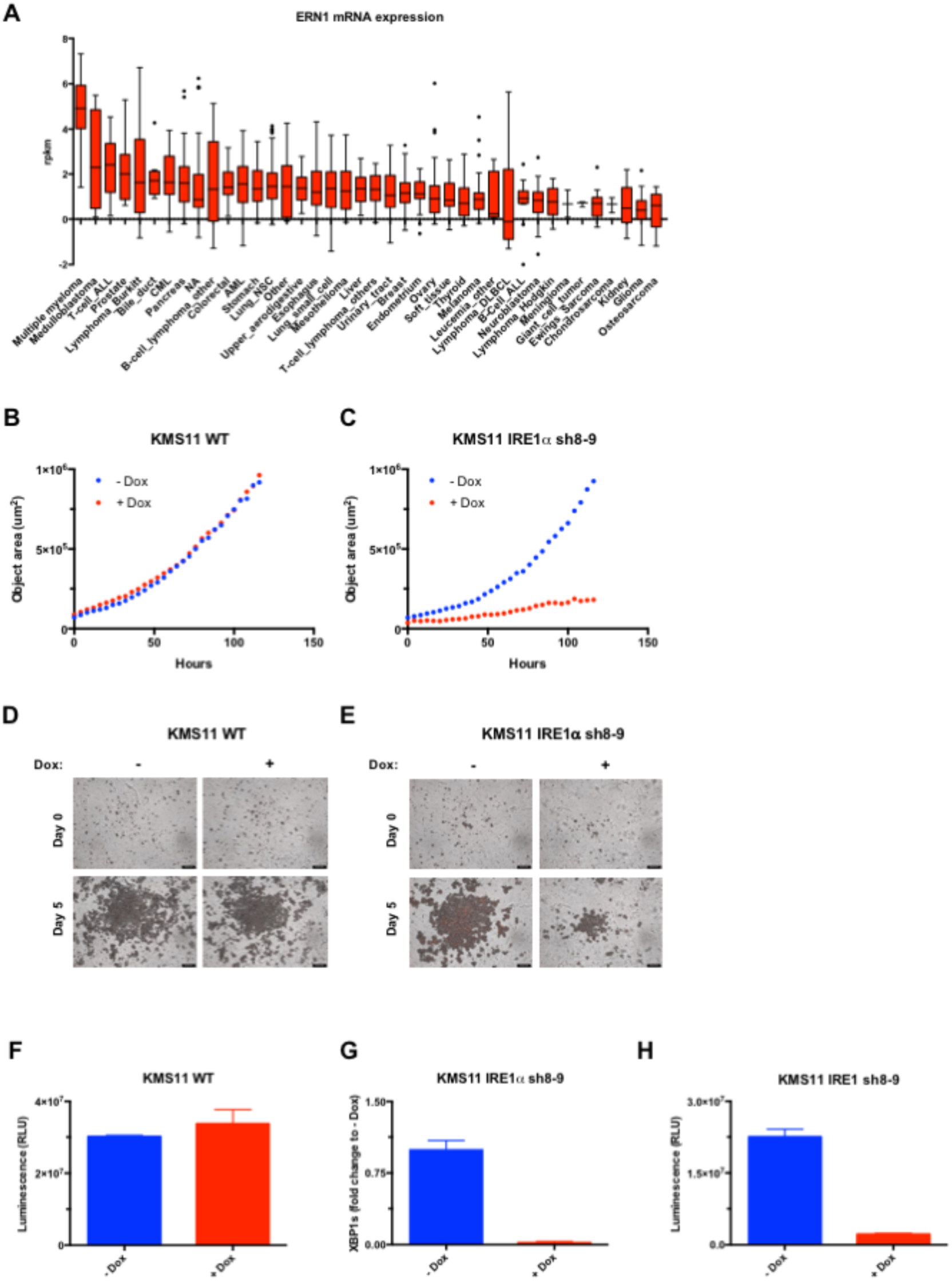
Expression of IREla in cancer cell lines and effect of its shRNA-based depletion on 3D spheroid growth of MM cells. (***A***) The cancer cell line encyclopedia (CCLE) dataset, which comprises RNAseq-based analysis of human cancer cell lines including 29 MM lines (Broad Institute, Cambridge, MA, USA) (https://portals.broadinstitute.org/ccle/page?gene=ERN1) was queried for expression of IRE1α (ERN1). (***B*** and ***C***) Non-transfected KMS11 cells (WT) or cells stably transfected with plasmids encoding Dox-inducible shRNAs against IRE1α were treated with Dox (0.5 μg/ml) for 3 days, seeded on Matrigel, allowed to grow as multiple 3D-spheroids, and analyzed over 5 days in an Incucyte™ instrument. (***F-H***) Cells were treated as in ***B*** and ***C*** and analyzed by CellTiterGlo^®^ assay to determine viability (***F-H***) or RT-QPCR to determine XBP1s mRNA levels (***G***).

**Fig S2.**
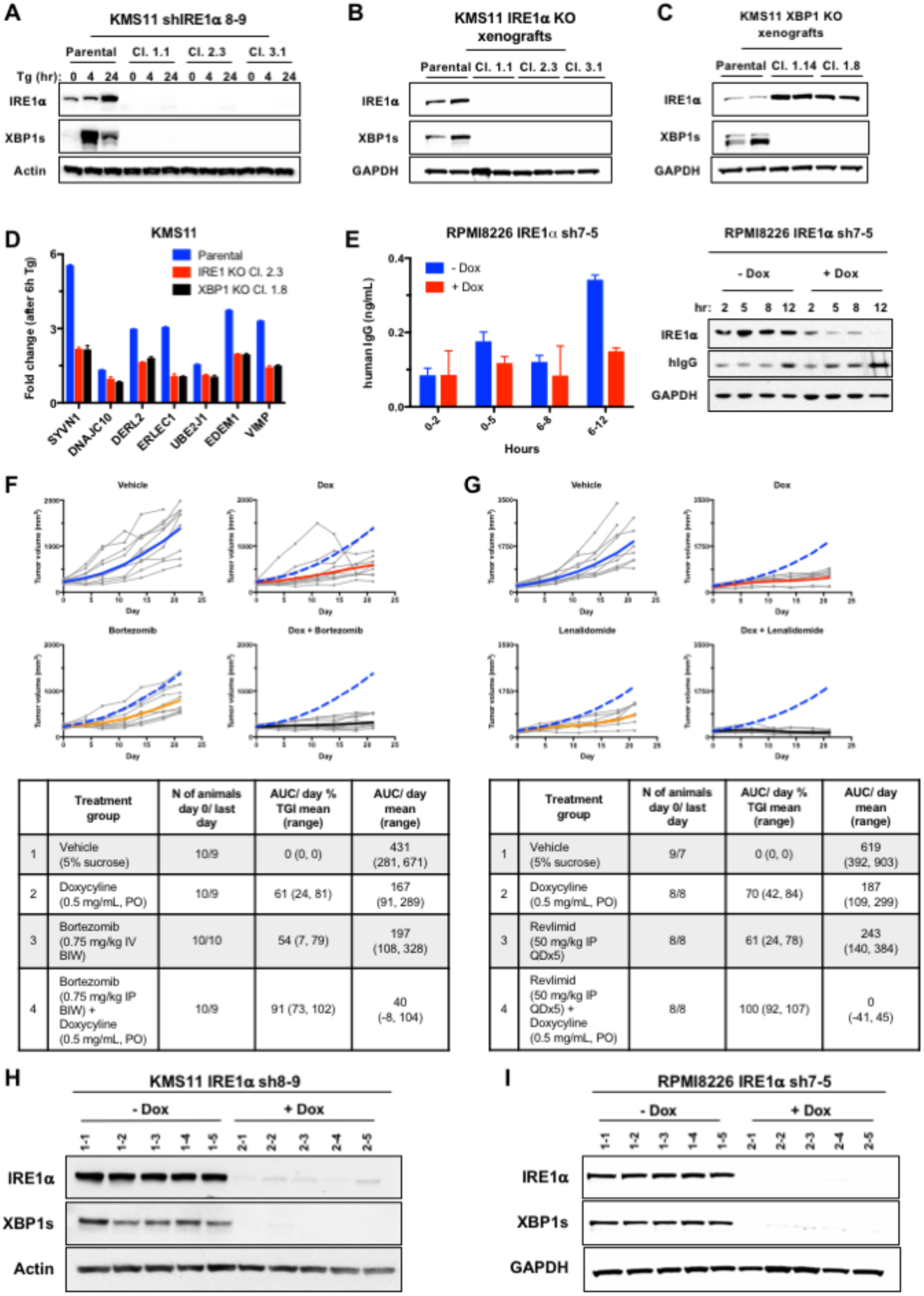
Genetic disruption of IRE1α attenuates secretory function and growth of subcutaneous human MM tumor xenografts in mice. (***A***) Parental and IRE1α KO KMS11 cells were treated with Tg (Tg, 100 nM) for the indicated time and analyzed by IB for expression of IRE1α and XBP1s protein. (***B*** and ***C***) IB analysis to confirm depletion of IRE1α and XBP1s in corresponding WT and KO KMS11 tumor xenografts. (D) IRE1α WT and KO cells were treated with Tg (100 nM) for 8 hr and analyzed by RNAseq. Fold-change in mRNA expression is shown for the indicated components of the ERAD machinery. (***E***) KMS11 cells stably transfected with a plasmid encoding shRNAs against IRE1α were incubated in the absence or presence of Dox (1 μg/ml) for up to 12 days. Levels of human IgG were analyzed in cell supernatants by ELISA (left-hand panel) or in cell lysates by IB (right-hand panel). (***F*** and ***G***) Tumor growth trajectories of individual animals, corresponding to the mean tumor volumes depicted in **Fig. *2D*** (***F***) and **Fig. 2E** (***G***). (***H*** and ***I***) IB analysis to confirming Dox-induced shRNA depletion of IRE1α and XBP1s in individual tumor xenografts.

**Figure S3.**
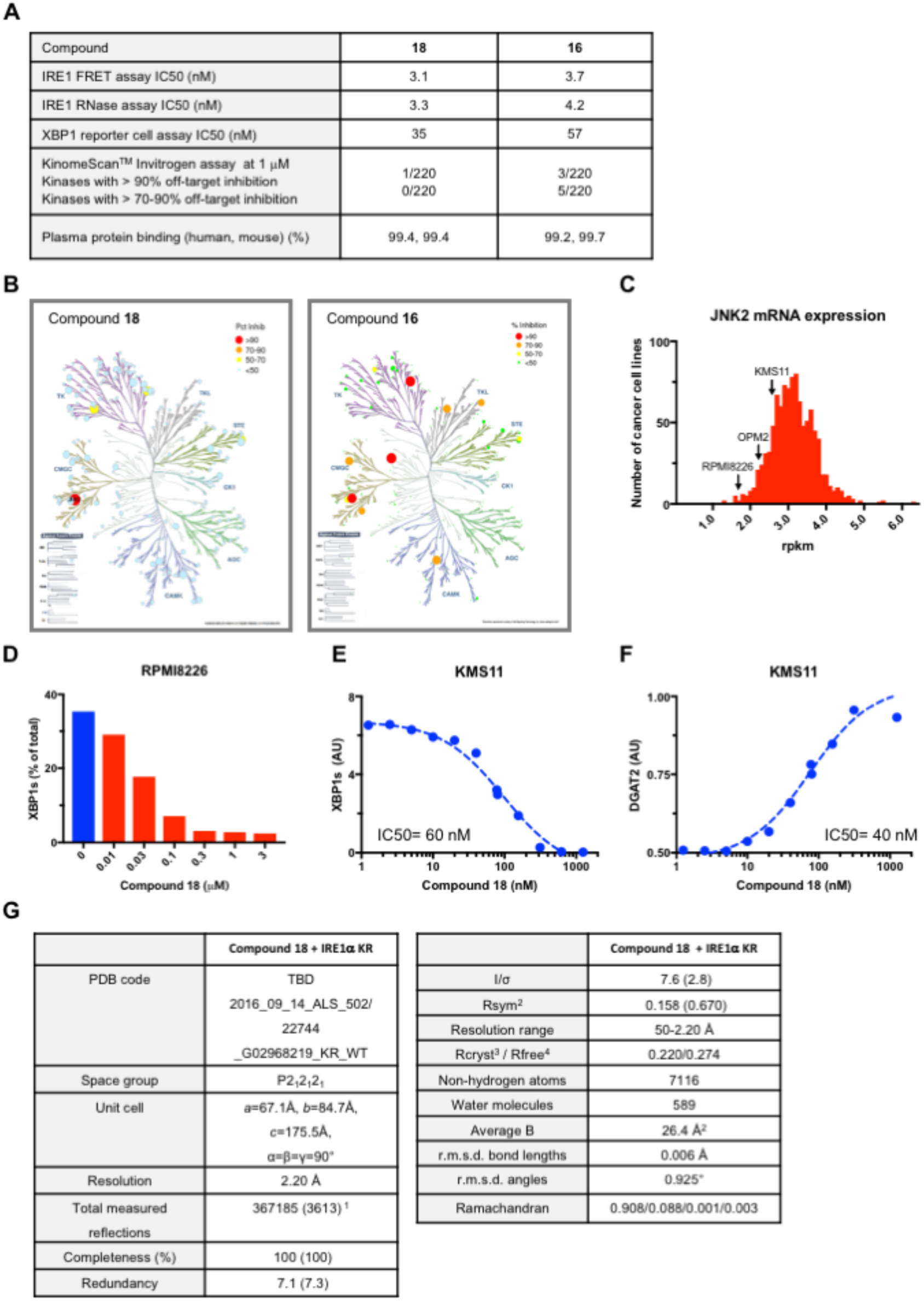
Biochemical characterization of compounds 18 and 16. (***A***) Biochemical properties and kinase selectivity of compound **18** and **16**. Kinase inhibition was determined by competition for binding of a staurosporine-based probe to the kinase pocket of a recombinant IRE1α protein comprising the kinase and endoribunuclease moieties. RNase activity of the recombinant protein was measured by cleavage of an XBP1s-based stem-loop structured RNA substrate as previously described (5). XBP1s cell reporter assay was performed as previously described (3, 5) (see Methods). Kinase-selectivity analysis against a panel of 220 kinases was performed at a compound concentration of 1 μM by KinomeScan™. (***B***) Schematic representation of the kinase interactions of **18** and **16**. Size and color of circles are related to interaction strength, as indicated in the top right inset. (***C***) Histogram of relative JNK2 mRNA expression in 1019 cancer cell lines in the CCLE dataset (Broad Institute, Cambridge, MA, USA). (***D***) RPMI8226 cells were incubated in the absence or presence of serial dilutions of **18** or **16**, for 8 hr and analyzed for constitutive XBP1s and XBP1u mRNA levels by RT-QPCR (%XBP1s mRNA is the ratio of XBP1s mRNA/(XBP1s mRNA+XBP1u mRNA). (E and ***F***) KMS11 cells were incubated for 8 hr with Tg (100 nM) in the absence or presence of **18** serial dilutions and analyzed by RT-QPCR for mRNA levels of XBP1s (***G***) or the RIDD target DGAT2 (***F***). Data in ***D-F*** are shown as the mean of triplicate determinations. (***G***) Crystallographic statistics for the structure of **18** with IRE1α.

**Fig S4.**
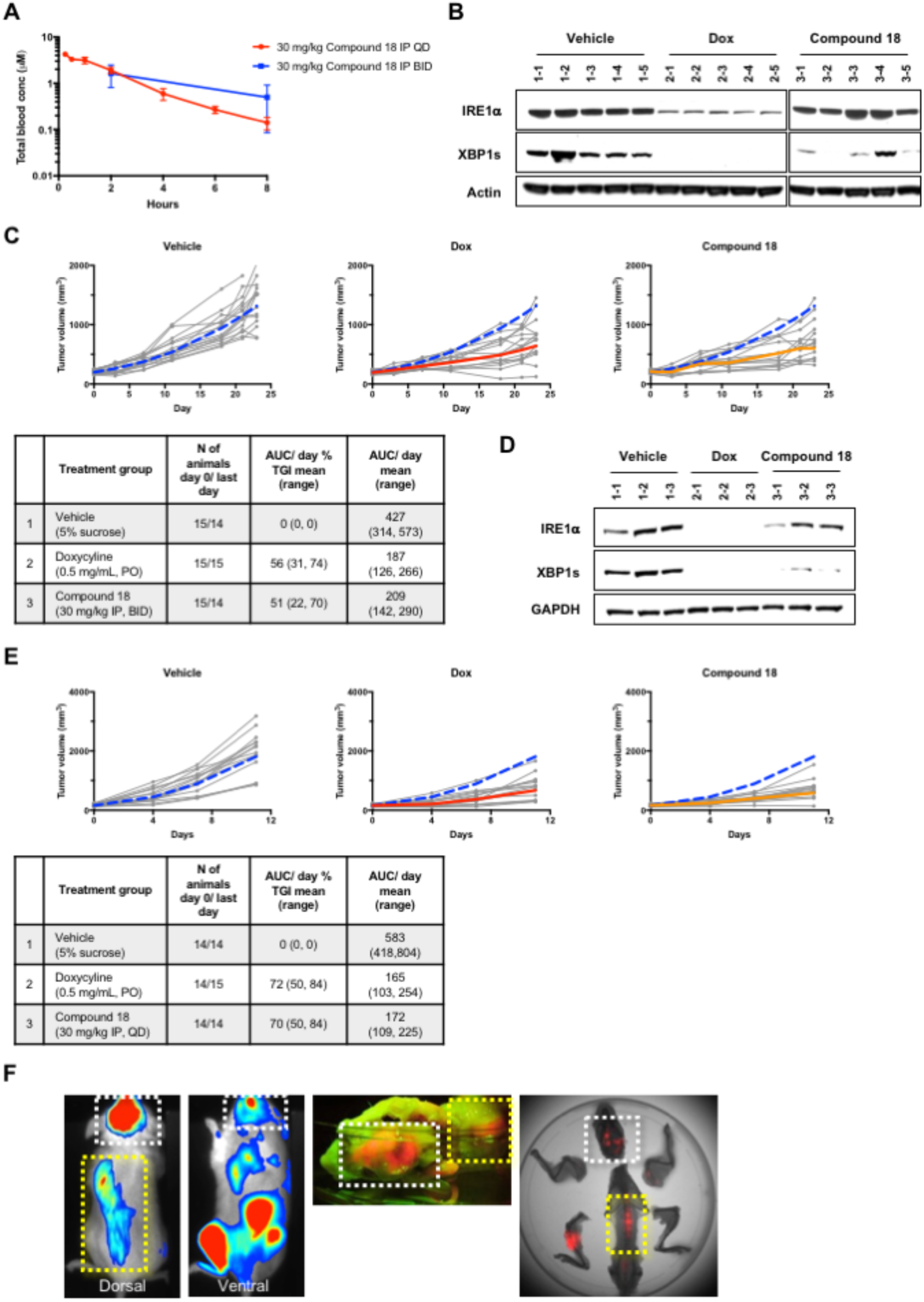
Small-molecule inhibition of IRE1α kinase attenuates XBP1s production and subcutaneous growth of human MM xenografts in mice. (***A***) C.B17-SCID mice bearing subcutaneous KMS11 tumor xenografts (3 per group) were treated either daily (QD) or twice per day (BID) intraperitoneally with **18** (30 mg/kg) over 4 days. Plasma was collected at the indicated time after the last dose and compound concentrations were determined by liquid chromatography and mass spectrometry. (***B***) Confirmation of XBP1s depletion in tumor xenografts sampled from individual mice depicted in **Fig. 4*A***. (***C***) Tumor growth trajectories of individual animals, corresponding to the mean tumor volumes depicted in **Fig. 4*A***. (***D***) Confirmation of XBP1s depletion in tumor xenografts sampled from individual mice depicted in **Fig. 4*B***. (***E***) Tumor growth trajectories of individual animals, corresponding to the mean tumor volumes depicted in **Fig. 4*B***. (***F***) RPMI8226 cells expressing plasmids encoding mCherry and luciferase were intravenously injected via the tail vein. Multifocal orthometastatic growth in the bone marrow (with typical skeletal lesions in the skull (white dashed box) and spine (yellow dashed box)) was confirmed by live bioluminescence (left-hand panels), post-mortal fluorescence imaging (middle panel), or fluorescence co-registered with X-ray imaging (right-hand panel) within same animals.

**Fig S5.**
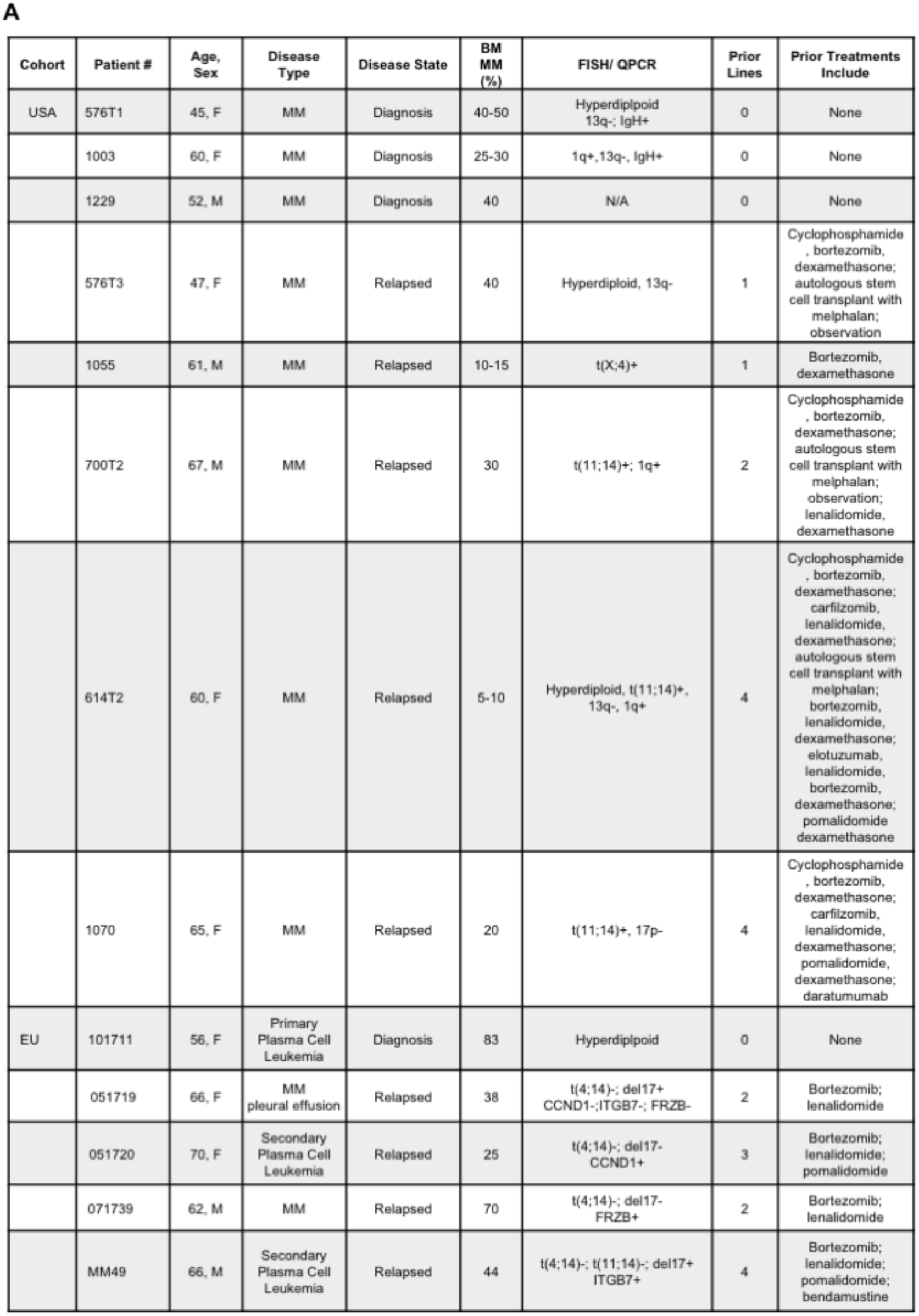
(***A***) Demographic, cytogenetic and treatment characteristics of bone marrow or peripheral blood samples donated by MM patient cohorts in the USA and EU as depicted in **Fig. 5**. HD, hyperdiploid. ASCT, autologous bone marrow transplant with melphalan.

**Fig S6.**
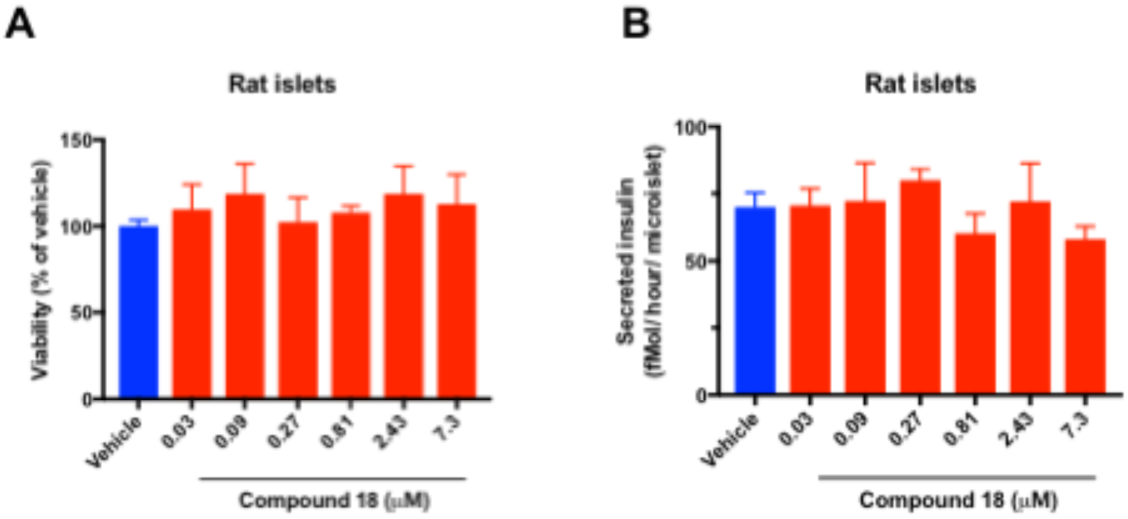
IRE1α kinase inhibition preserves survival and insulin secretion by pancreatic islet 3D microtissues. Rat pancreatic islets were isolated, dissociated into single cells, replated in microtiter wells (1000 cells/drop), and allowed to form 3D microtissues of ~120 μM in diameter over 7 days using inSphero™ technology. Microtissues (n=5 per treatment) were then incubated for 7 days in the absence or presence of compound **18** at the indicated concentrations, and then (***A***) analyzed for cell viability by ATP levels; or (***B***) challenged with glucose (16.7 mM) for 1 hr and analyzed for insulin secretion by ELISA.

## Notes

#### Summary of Updates

Minor typo - author list

