## Supplementary material for "Disruption of IRE1α through its Kinase Domain Attenuates Multiple Myeloma"

**This PDF file includes:**

- Supplemental Material and Methods
- Figs. S1 to S6
- References for SI reference citations

### Supplemental Material and Methods

**Generation of IRE1α shRNA-expressing cells.** Cells were infected with shRNAs by lentivirus using the pG-GW-pHUSH vector system. Briefly $1x10^7$ 293T cells were plated in 6-well dishes, allowed to grow for 24 hr and transfected with lentivirus plasmids using Lipofectamine 2000 (Invitrogen) following the manufacturer's protocol. After 6 hr the media was replaced, and after another 24 hr the virus was harvested from the cells and filtered through a 0.45 mm tube-top filter. Cleared supernatants containing the viral particles and 8 mg/ml polybreen were added to the target-cell wells, spun at 1800 rpm for 45 min at room temperature, and placed back into the incubator. After 48 hr cells were subjected to selection with with puromycin containing media. After 10 passages, cells were tested for virus by Lentivirus X p24 Rapid Titer Kit (Clonetech) then sorted using FACS and RFP+ selection.

IRE1α shRNA sequences (antisense):

shRNA5:
AGCTTTTCCAAAAAACCAAGATGCTGGAGAGATTTCTCTTGATAATCTCTCCA
GCATCTTGGGGG

shRNA7:
AGCTTTTCCAAAAAAAGAACAAGCTCAACTACTTTCTCTTGATAAGTAGTTGA
GCTTGTTCTGGG

shRNA8:
AGCTTTTCCAAAAAAGCACGTGAATTGATAGAGATCTCTTGAATCTCTATCAA
TTCACGTGCGGG

shRNA9:
AGCTTTTCCAAAAAAGAGAAGATGATTGCGATGGTCTCTTGAACCATCGCAA
TCATCTTCTCGGG

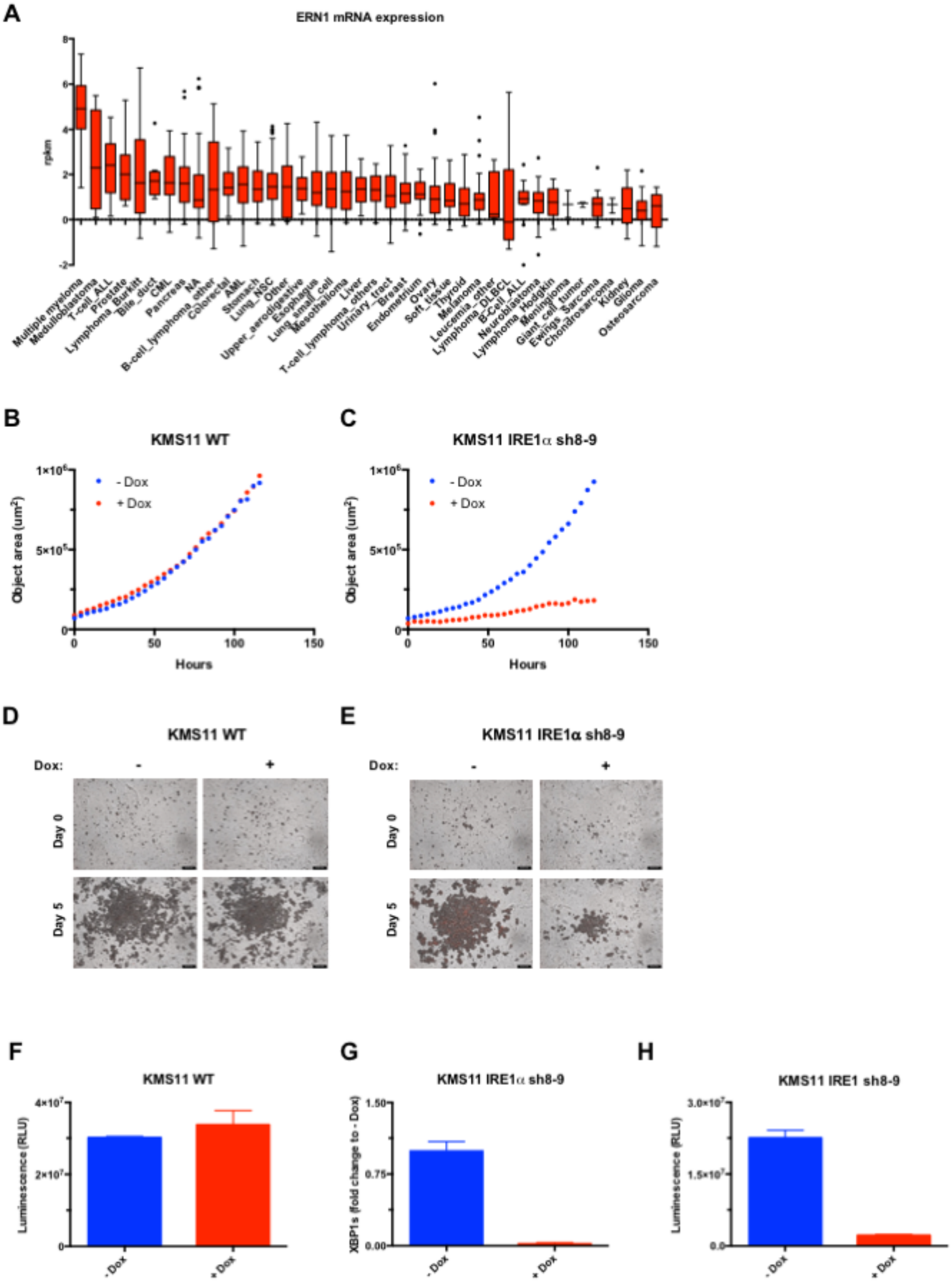


**Fig. S1**. **Expression of IRE1α in cancer cell lines and effect of its shRNA-based depletion on 3D spheroid growth of MM cells.** (*A*) The cancer cell line encyclopedia (CCLE) dataset, which comprises RNAseq-based analysis of human cancer cell lines including 29 MM lines (Broad Institute, Cambridge, MA, USA) (https://portals.broadinstitute.org/ccle/page?gene=ERN1) was queried for expression of IRE1α (ERN1). (***B*** and ***C***) Non-transfected KMS11 cells (WT) or cells stably transfected with plasmids encoding Dox-inducible shRNAs against IRE1α were treated with Dox (0.5 μg/ml) for 3 days, seeded on Matrigel, allowed to grow as multiple 3D-spheroids, and analyzed over 5 days in an Incucyte™ instrument. (***F***-***H***) Cells were treated as in ***B*** and ***C*** and analyzed by CellTiterGlo® assay to determine viability (***F***-***H***) or RT-QPCR to determine XBP1s mRNA levels (***G***).

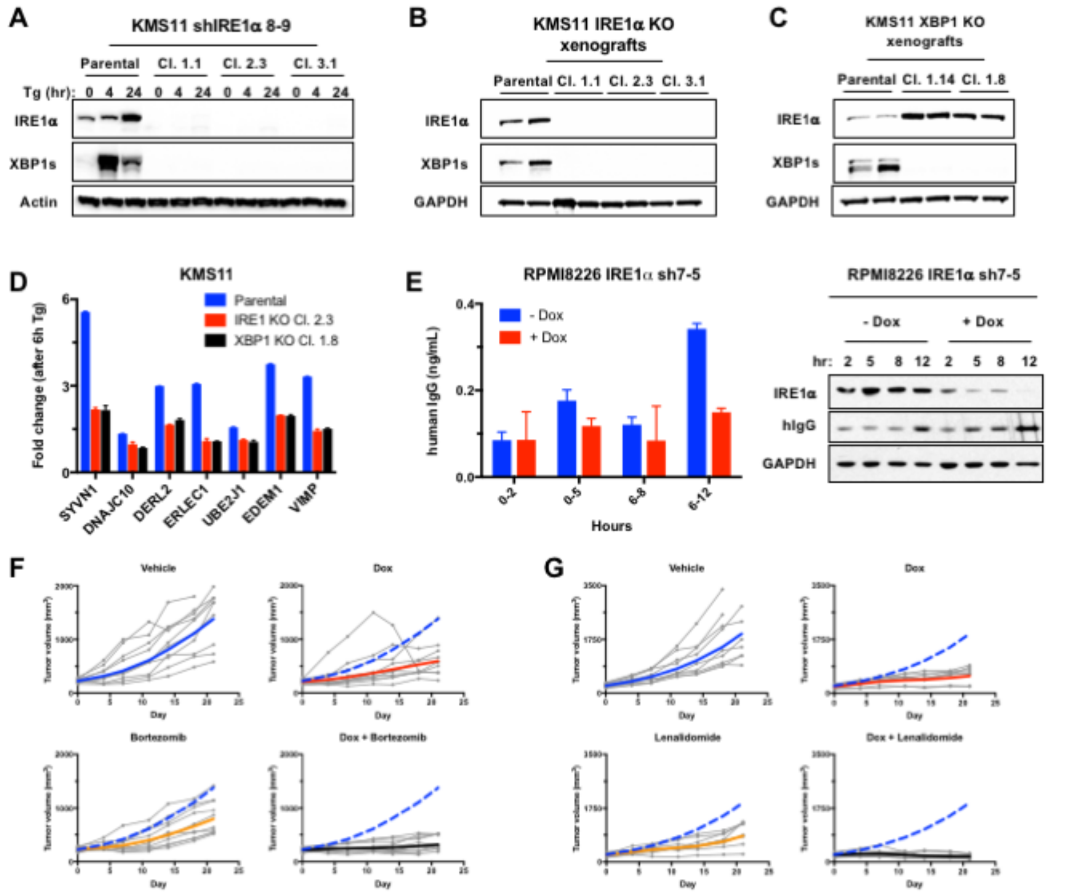


| | Treatment group | N of animals day 0/ last day | AUC/ day % TGI mean (range) | AUC/ day mean (range) |
|---|---|---|---|---|
| 1 | Vehicle (5% sucrose) | 10/9 | 0 (0, 0) | 431 (281, 671) |
| 2 | Doxycyline (0.5 mg/mL, PO) | 10/9 | 61 (24, 81) | 167 (91, 289) |
| 3 | Bortezomib (0.75 mg/kg IV BIW) | 10/10 | 54 (7, 79) | 197 (108, 328) |
| 4 | Bortezomib (0.75 mg/kg IP BIW) + Doxycyline (0.5 mg/mL, PO) | 10/9 | 91 (73, 102) | 40 (-8, 104) |

| | Treatment group | N of animals day 0/ last day | AUC/ day % TGI mean (range) | AUC/ day mean (range) |
|---|---|---|---|---|
| 1 | Vehicle (5% sucrose) | 9/7 | 0 (0, 0) | 619 (392, 903) |
| 2 | Doxycyline (0.5 mg/mL, PO) | 8/8 | 70 (42, 84) | 187 (109, 299) |
| 3 | Revlimid (50 mg/kg IP QDx5) | 8/8 | 61 (24, 78) | 243 (140, 384) |
| 4 | Revlimid (50 mg/kg IP QDx5) + Doxycyline (0.5 mg/mL, PO) | 8/8 | 100 (92, 107) | 0 (-41, 45) |

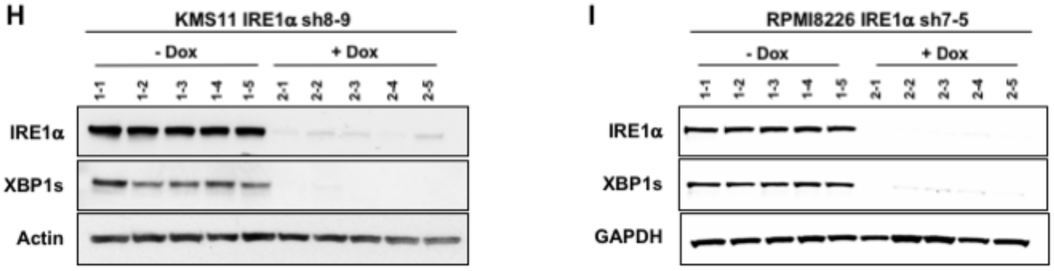


**Fig. S2. Genetic disruption of IRE1α attenuates secretory function and growth of subcutaneous human MM tumor xenografts in mice.** (*A*) Parental and IRE1α KO

KMS11 cells were treated with Tg (Tg, 100 nM) for the indicated time and analyzed by IB for expression of IRE1α and XBP1s protein. (***B*** and ***C***) IB analysis to confirm depletion of IRE1α and XBP1s in corresponding WT and KO KMS11 tumor xenografts. (***D***) IRE1α WT and KO cells were treated with Tg (100 nM) for 8 hr and analyzed by RNAseq. Fold-change in mRNA expression is shown for the indicated components of the ERAD machinery. (***E***) KMS11 cells stably transfected with a plasmid encoding shRNAs against IRE1α were incubated in the absence or presence of Dox (1 μg/ml) for up to 12 days. Levels of human IgG were analyzed in cell supernatants by ELISA (left-hand panel) or in cell lysates by IB (right-hand panel). (***F*** and ***G***) Tumor growth trajectories of individual animals, corresponding to the mean tumor volumes depicted in **Fig. 2*D*** (***F***) and **Fig. 2*E*** (***G***). (***H*** and ***I***) IB analysis to confirming Dox-induced shRNA depletion of IRE1α and XBP1s in individual tumor xenografts.

**A**

| Compound | 18 | 16 |
|---|---|---|
| IRE1 FRET assay IC50 (nM) | 3.1 | 3.7 |
| IRE1 RNase assay IC50 (nM) | 3.3 | 4.2 |
| XBP1 reporter cell assay IC50 (nM) | 35 | 57 |
| KinomeScan™ Invitrogen assay at 1 µM<br>Kinases with > 90% off-target inhibition<br>Kinases with > 70-90% off-target inhibition | 1/220<br>0/220 | 3/220<br>5/220 |
| Plasma protein binding (human, mouse) (%) | 99.4, 99.4 | 99.2, 99.7 |

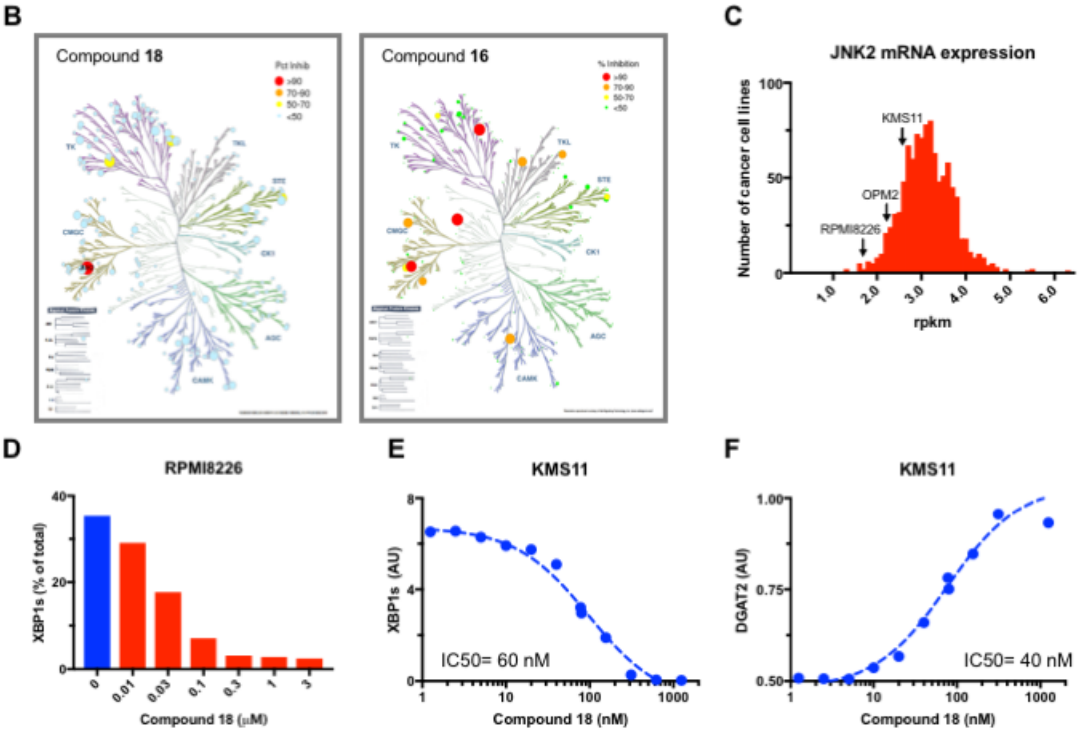


**G**

| | Compound 18 + IRE1α KR |
|---|---|
| PDB code | TBD<br>2016_09_14_ALS_502/<br>22744<br>_G02968219_KR_WT |
| Space group | $P2_12_12_1$ |
| Unit cell | *a*=67.1Å, *b*=84.7Å,<br>*c*=175.5Å,<br>α=β=γ=90° |
| Resolution | 2.20 Å |
| Total measured reflections | 367185 (3613)[1] |
| Completeness (%) | 100 (100) |
| Redundancy | 7.1 (7.3) |

| | Compound 18 + IRE1α KR |
|---|---|
| I/σ | 7.6 (2.8) |
| Rsym[2] | 0.158 (0.670) |
| Resolution range | 50-2.20 Å |
| Rcryst[3] / Rfree[4] | 0.220/0.274 |
| Non-hydrogen atoms | 7116 |
| Water molecules | 589 |
| Average B | 26.4 Å$^2$ |
| r.m.s.d. bond lengths | 0.006 Å |
| r.m.s.d. angles | 0.925° |
| Ramachandran | 0.908/0.088/0.001/0.003 |

**Figure S3**. **Biochemical characterization of compounds 18 and 16.** (*A*) Biochemical properties and kinase selectivity of compound **18** and **16**. Kinase inhibition was determined by competition for binding of a staurosporine-based probe to the kinase pocket of a recombinant IRE1α protein comprising the kinase and endoribunuclease moieties. RNase activity of the recombinant protein was measured by cleavage of an XBP1s-based stem-loop structured RNA substrate as previously described (5). XBP1s cell reporter assay was performed as previously described (3, 5) (see Methods). Kinase-selectivity analysis against a panel of 220 kinases was performed at a compound concentration of 1 μM by KinomeScan™. (***B***) Schematic representation of the kinase interactions of **18** and **16**. Size and color of circles are related to interaction strength, as indicated in the top right inset. (***C***) Histogram of relative JNK2 mRNA expression in 1019 cancer cell lines in the CCLE dataset (Broad Institute, Cambridge, MA, USA). (***D***) RPMI8226 cells were incubated in the absence or presence of serial dilutions of **18** or **16**, for 8 hr and analyzed for constitutive XBP1s and XBP1u mRNA levels by RT-QPCR (%XBP1s mRNA is the ratio of XBP1s mRNA/(XBP1s mRNA+XBP1u mRNA). (***E*** and ***F***) KMS11 cells were incubated for 8 hr with Tg (100 nM) in the absence or presence of **18** serial dilutions and analyzed by RT-QPCR for mRNA levels of XBP1s (***G***) or the RIDD target DGAT2 (***F***). Data in ***D-F*** are shown as the mean of triplicate determinations. (***G***) Crystallographic statistics for the structure of **18** with IRE1α.

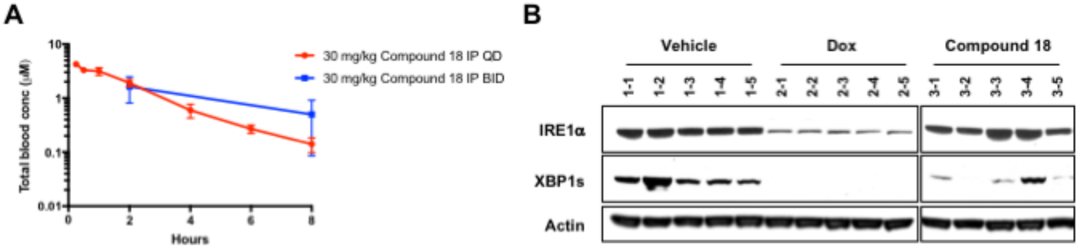


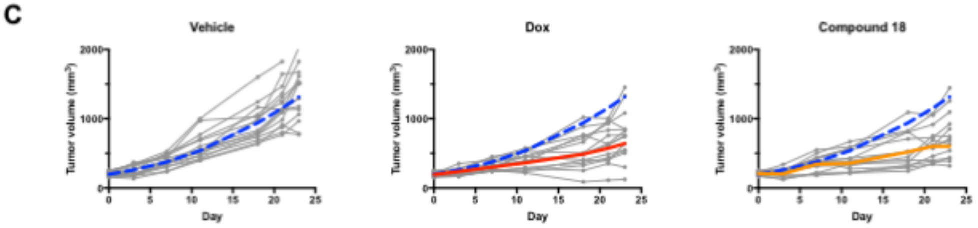


| | Treatment group | N of animals day 0/ last day | AUC/ day % TGI mean (range) | AUC/ day mean (range) |
|---|---|---|---|---|
| 1 | Vehicle (5% sucrose) | 15/14 | 0 (0, 0) | 427 (314, 573) |
| 2 | Doxycyline (0.5 mg/mL, PO) | 15/15 | 56 (31, 74) | 187 (126, 266) |
| 3 | Compound 18 (30 mg/kg IP, BID) | 15/14 | 51 (22, 70) | 209 (142, 290) |

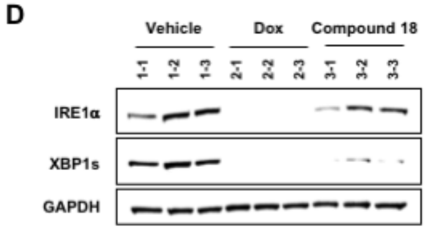


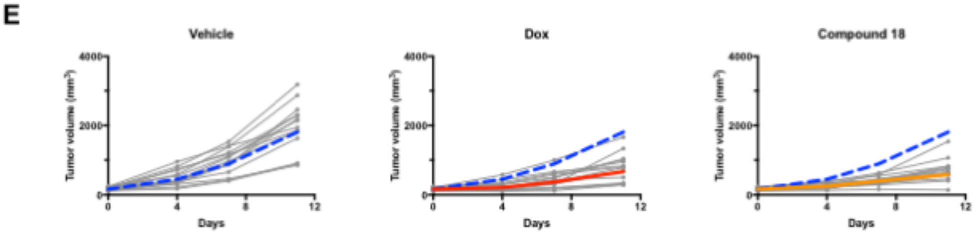


| | Treatment group | N of animals day 0/ last day | AUC/ day % TGI mean (range) | AUC/ day mean (range) |
|---|---|---|---|---|
| 1 | Vehicle (5% sucrose) | 14/14 | 0 (0, 0) | 583 (418,804) |
| 2 | Doxycyline (0.5 mg/mL, PO) | 14/15 | 72 (50, 84) | 165 (103, 254) |
| 3 | Compound 18 (30 mg/kg IP, QD) | 14/14 | 70 (50, 84) | 172 (109, 225) |

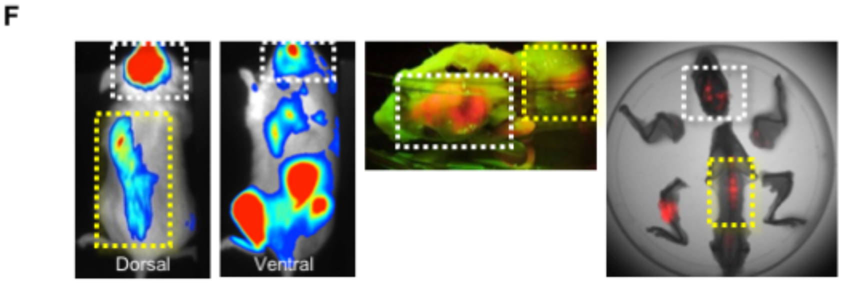


**Fig. S4**. **Small-molecule inhibition of IRE1α kinase attenuates XBP1s production and subcutaneous growth of human MM xenografts in mice.** (*A*) C.B17-SCID mice

bearing subcutaneous KMS11 tumor xenografts (3 per group) were treated either daily (QD) or twice per day (BID) intraperitoneally with **18** (30 mg/kg) over 4 days. Plasma was collected at the indicated time after the last dose and compound concentrations were determined by liquid chromatography and mass spectrometry. (***B***) Confirmation of XBP1s depletion in tumor xenografts sampled from individual mice depicted in **Fig. 4*A***. (***C***) Tumor growth trajectories of individual animals, corresponding to the mean tumor volumes depicted in **Fig. 4*A***. (***D***) Confirmation of XBP1s depletion in tumor xenografts sampled from individual mice depicted in **Fig. 4*B***. (***E***) Tumor growth trajectories of individual animals, corresponding to the mean tumor volumes depicted in **Fig. 4*B***.
(***F***) RPMI8226 cells expressing plasmids encoding mCherry and luciferase were intravenously injected via the tail vein. Multifocal orthometastatic growth in the bone marrow (with typical skeletal lesions in the skull (white dashed box) and spine (yellow dashed box)) was confirmed by live bioluminescence (left-hand panels), post-mortal fluorescence imaging (middle panel), or fluorescence co-registered with X-ray imaging (right-hand panel) within same animals.

**A**

| Cohort | Patient # | Age, Sex | Disease Type | Disease State | BM MM (%) | FISH/ QPCR | Prior Lines | Prior Treatments Include |
|---|---|---|---|---|---|---|---|---|
| USA | 576T1 | 45, F | MM | Diagnosis | 40-50 | Hyperdiplpoid 13q-; IgH+ | 0 | None |
| | 1003 | 60, F | MM | Diagnosis | 25-30 | 1q+,13q-, IgH+ | 0 | None |
| | 1229 | 52, M | MM | Diagnosis | 40 | N/A | 0 | None |
| | 576T3 | 47, F | MM | Relapsed | 40 | Hyperdiploid, 13q- | 1 | Cyclophosphamide , bortezomib, dexamethasone; autologous stem cell transplant with melphalan; observation |
| | 1055 | 61, M | MM | Relapsed | 10-15 | t(X;4)+ | 1 | Bortezomib, dexamethasone |
| | 700T2 | 67, M | MM | Relapsed | 30 | t(11;14)+; 1q+ | 2 | Cyclophosphamide , bortezomib, dexamethasone; autologous stem cell transplant with melphalan; observation; lenalidomide, dexamethasone |
| | 614T2 | 60, F | MM | Relapsed | 5-10 | Hyperdiploid, t(11;14)+, 13q-, 1q+ | 4 | Cyclophosphamide , bortezomib, dexamethasone; carfilzomib, lenalidomide, dexamethasone; autologous stem cell transplant with melphalan; bortezomib, lenalidomide, dexamethasone; elotuzumab, lenalidomide, bortezomib, dexamethasone; pomalidomide dexamethasone |
| | 1070 | 65, F | MM | Relapsed | 20 | t(11;14)+, 17p- | 4 | Cyclophosphamide , bortezomib, dexamethasone; carfilzomib, lenalidomide, dexamethasone; pomalidomide, dexamethasone; daratumumab |
| EU | 101711 | 56, F | Primary Plasma Cell Leukemia | Diagnosis | 83 | Hyperdiplpoid | 0 | None |
| | 051719 | 66, F | MM pleural effusion | Relapsed | 38 | t(4;14)-; del17+ CCND1-;ITGB7-; FRZB- | 2 | Bortezomib; lenalidomide |
| | 051720 | 70, F | Secondary Plasma Cell Leukemia | Relapsed | 25 | t(4;14)-; del17- CCND1+ | 3 | Bortezomib; lenalidomide; pomalidomide |
| | 071739 | 62, M | MM | Relapsed | 70 | t(4;14)-; del17- FRZB+ | 2 | Bortezomib; lenalidomide |
| | MM49 | 66, M | Secondary Plasma Cell Leukemia | Relapsed | 44 | t(4;14)-; t(11;14)-; del17+ ITGB7+ | 4 | Bortezomib; lenalidomide; pomalidomide; bendamustine |

**Fig. S5**. (***A***) Demographic, cytogenetic and treatment characteristics of bone marrow or peripheral blood samples donated by MM patient cohorts in the USA and EU as depicted in **Fig. 5**. HD, hyperdiploid. ASCT, autologous bone marrow transplant with melphalan.

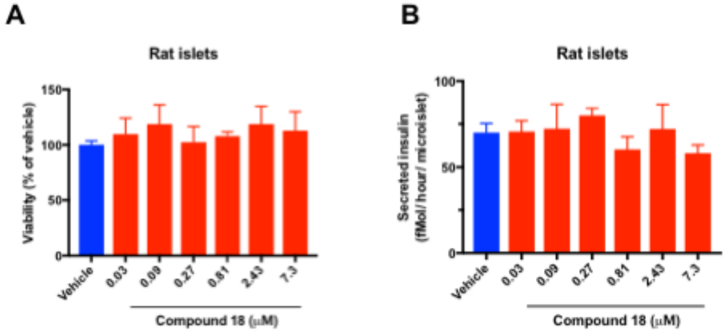


**Fig. S6**. **IRE1α kinase inhibition preserves survival and insulin secretion by pancreatic islet 3D microtissues.** Rat pancreatic islets were isolated, dissociated into single cells, replated in microtiter wells (1000 cells/drop), and allowed to form 3D microtissues of ~120 μM in diameter over 7 days using inSphero™ technology. Microtissues (n=5 per treatment) were then incubated for 7 days in the absence or presence of compound **18** at the indicated concentrations, and then (***A***) analyzed for cell viability by ATP levels; or (***B***) challenged with glucose (16.7 mM) for 1 hr and analyzed for insulin secretion by ELISA.
